# Input-space geometry shapes adaptation dynamics underlying repetition suppression in a neural network model of relatedness priming

**DOI:** 10.64898/2026.09.13.751213

**Authors:** Diletta Bartolini, Thomas P. Reber, Tatjana Tchumatchenko, Matthias Voigt

**Affiliations:** Department of Mathematics, University of Pavia, Pavia, Italy; Faculty of Mathematics and Computer Science, UniDistance Suisse, Brig, Switzerland; Faculty of Psychology, UniDistance Suisse, Brig, Switzerland; Department of Neuroscience, Medical Faculty, University of Bonn, Germany; Institute of Experimental Epileptology and Cognition Research, Life and Brain Center, Universitätsklinikum Bonn, Germany

**Keywords:** Neural adaptation, adaptive exponential integrate-and-fire model, triplet spike-timing-dependent plasticity, sharpening, fatiguing

## Abstract

A key function of the human brain is its ability to dynamically adapt to novel contexts by integrating prior experience. While neural adaptation is widely observed across cortical systems, the microcircuit-level mechanisms governing its intensity and temporal dynamics remain unclear. To bridge this gap, we develop a recurrent network of adaptive exponential integrate-and-fire neurons governed by a triplet spike-timing-dependent plasticity rule, designed to reproduce neural dynamics recorded via intracranial electrophysiology and single-unit recordings from the medial temporal lobe of neurosurgical patients performing a priming task. Using input organizations inspired by the experimental paradigm, we systematically vary input geometry to investigate its impact on adaptation dynamics. We find that neural adaptation emerges from recurrent dynamics shaped by learned connectivity, with both its magnitude and temporal profile depending on the geometry of the input space. This dependence gives rise, at the simulated single-neuron level, to a continuum of response regimes ranging from sharpening-like to fatiguing-like dynamics. Sharpening-like responses dominate when inputs exhibit high within-meta-category similarity and strong between-meta-category separation, whereas fatiguing-like responses emerge under the opposite regime.

## 1 Introduction

Neural adaptation is a fundamental property of cortical circuits, reflecting the brain’s capacity to optimize its responses based on recent experience [36]. It can manifest as priming, a facilitation effect in which responses to repeated (repetition priming) or semantically related (relatedness priming) stimuli become faster and more accurate [19].

The underlying neuronal mechanisms of adaptation and the resulting behavioral optimization are still being discussed [19]. Among the most influential formulations are spreading activation models [26], in which semantic knowledge is represented as a network of interconnected nodes, each corresponding to a concept or semantic feature. Activation of one node propagates to its neighbors, increasing their pre-existing activity through residual activation spread, so that target-evoked activity is added to this elevated baseline, making threshold-crossing more likely even though the threshold itself remains constant, thereby facilitating subsequent activation of related concepts. Within this framework, behavioral facilitation naturally emerges from the dynamic propagation of activation across the network. While highly influential in linguistic and cognitive domains [10], these models remain largely abstract, operating at a symbolic rather than mechanistic level.

Only a few studies attempt to incorporate biological constraints, such as neuromodulatory influences on activation spread [3], yet they provide limited insight into how large-scale behavioral facilitation arises from underlying neural mechanisms. In particular, they do not explain how neural adaptation observed at mesoscopic and macroscopic scales, as revealed by neuroimaging studies [7, 20, 35], emerges from the dynamics of interacting neuronal populations. Despite extensive empirical characterization of adaptation phenomena, its underlying microcircuit mechanisms remain poorly understood. Bridging this explanatory gap requires mechanistic models that are both biologically plausible and capable of reproducing the temporal and intensity profiles of adaptation observed experimentally.

Among the available modeling frameworks, the adaptive exponential integrate-and-fire (AdEx) model [6] has emerged as a particularly promising choice. Building on the classical leaky integrate-and-fire model, it incorporates an exponential term that captures the rapid voltage rise during spike initiation alongside an adaptation variable that accumulates with neuronal activity and exerts a subtractive, hyperpolarizing influence on the membrane potential, thereby reducing excitability without altering the spike threshold itself.

To ground this mechanistic model in experimental evidence, it is therefore important to rely on high-precision neurophysiological recordings, which serve as a quantitative benchmark for evaluating the model’s accuracy. Among available techniques, invasive electrophysiology offers access to neural dynamics across complementary spatial scales, from mesoscopic population activity to microscopic single-neuron activity, while preserving high temporal resolution [21]. Intracranial electrophysiology (iEEG), recorded from electrodes implanted both on the cortical surface and within deep brain structures such as the medial temporal lobe, captures mesoscale population signals that reflect the summed activity of large neuronal ensembles. Complementing this, microwire recordings provide access to single-neuron activity and true local field potentials at the microscale. Together, these modalities enable the investigation of the microcircuit mechanisms driving mesoscopic population activity and their contribution to large-scale cortical dynamics.

A particularly relevant study employing this approach is provided by *Reber et al*. in [33], who simultaneously recorded iEEG and single-unit activity across multiple medial temporal lobe regions in epileptic patients. Their results reveal a clear correspondence between behavioral and neural adaptation, observed at both single-neuron and population levels. At the macroscopic scale, iEEG event-related potentials show temporal compression following repeated or related stimuli, indicating more efficient and temporally focused neural processing.

Furthermore, they demonstrate that the reduction of event-related potentials in the macroscopic iEEG signal is associated with distinct single-neuron adaptation patterns across medial temporal lobe subregions. At the level of individual neurons, two commonly described adaptation phenotypes have been characterized. The first, termed sharpening [19, 42], describes a selective reduction in responses to non-preferred stimuli: priming effectively narrows a neuron’s tuning curve around its most effective stimulus, increasing response selectivity without uniformly attenuating activity. The second, termed fatiguing [13, 23], describes a more global reduction of responses across stimuli, largely independent of their relative efficacy for the recorded neuron. A third proposed mechanism, facilitation [19], predicts enhanced or faster responses upon repetition, although robust single-unit evidence for this mode in the human medial temporal lobe remains absent [33].

These profiles carry distinct mechanistic interpretations. Sharpening has been attributed to fast recurrent or inhibitory competition [27] that selectively suppresses responses to non-preferred stimuli on a sub-second timescale, effectively sparing the connections encoding diagnostic stimulus features [42]. Fatiguing, by contrast, has been linked to population-level adaptation processes, such as spike-frequency adaptation or short-term synaptic depression, that reduce excitability following repeated activation, with weaker dependence on stimulus identity [11, 13]. Both modes have been discussed within predictive coding frameworks [15], where repeated stimuli elicit reduced prediction errors; sharpening would correspond to more selective representational updating, whereas fatiguing would reflect a more uniform attenuation of prediction error signals. However, dissociating these mechanisms in experimental data remains challenging, as they often coexist within the same cortical region and may even be observed in adjacent neurons [11, 33]. Computational models therefore provide a principled framework to probe which circuit properties and input statistics favor one regime over the other.

Motivated by these findings, we develop a biologically grounded spiking network model to reproduce the dynamics reported in [33]. The network comprises two distinct populations, excitatory regular-spiking and inhibitory fast-spiking neurons, with the dynamics of each neuron modeled using the AdEx model. Synaptic weights are trained according to a triplet-based spike-timing-dependent plasticity rule [32], following the paradigm described in [16], and subsequently held fixed during all testing phases. This synaptic rule extends the classical pair-based formulation by incorporating the dependence of synaptic changes on the repetition frequency of spike pairs, providing a more biologically realistic mechanism for experience-dependent adaptation.

To enable a direct comparison with prior findings, we first evaluated the original behavioral paradigm using the same image dataset. We then extend this framework by employing synthetically generated input feature vectors, characterized by three parameters controlling stimulus similarity and categorical structure. This integrative approach allows us to systematically relate adaptation dynamics to the statistical structure of stimuli, providing a mechanistic framework for how recurrent cortical networks encode and adapt to structured sensory information.

## 2 Results

The network, fully described in the Methods section, is driven by two types of image inputs, each comprising *S* = 100 feature vectors distributed across two meta-categories, each further divided into five semantic categories.

The first input type consists of feature vectors encoded using the DINO vision transformer model [8], derived from real visual stimuli presented during the invasive electrophysiology recordings of [33]. The second is synthetic, parameterized by three variables (*α, β, ε*) and designed to approximate the structure of the image-derived features.

In this section, we present results from simulations driven by the synthetic input. A total of 200 distinct sequences of synthetic feature vectors (SFVs) are generated, each corresponding to a trial of five stimuli. SFVs are labeled individually according to two conditions: *primed*, when the current SFV belongs to the same semantic category as the immediately preceding one, and *control*, when it belongs to a different category. The first SFV of each sequence is excluded from categorization, as no preceding context is available, while the remaining SFVs are equally distributed across the two conditions. Sequences are generated randomly under the constraint of maintaining this balanced condition distribution. To ensure uniform coverage of the stimulus set and prevent the over-representation of any individual item, each SFV is removed from the pool upon presentation and only reintroduced once the entire set has been exhausted. Each simulation lasts 1.6 s. During the initial 100 ms, the network is driven by a stochastic sinusoidal input current, introduced to mimic spontaneous baseline activity and to prevent network quiescence [12], see the Methods section for details. Following the initialization phase, the network receives the sequence of SFVs for the trial, with each vector presented for 300 ms without inter-stimulus intervals.

Corresponding simulations with DINO-extracted feature vectors are reported in the DINO feature embeddings appendix.

### 2.1 Analysis of SFVs

To characterize the structure of the synthetic input, we compute the cosine similarity score between two feature vectors **f**_*i*_, **f**_*j*_ *∈* ℝ^*N*^ (with *N* = 768, see External input current), defined as

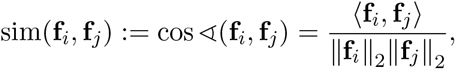

where ⟨·,· ⟩, and ‖ · ‖ _2_ denote the standard Euclidean inner product and its induced norm, respectively. Values close to 1 indicate strong alignment, whereas lower values indicate weaker similarity.

Applied to the SFVs input, this analysis reveals a highly structured representational space (Figure 1a). SFVs within the same category exhibit similarity values approaching 1, indicating nearly identical representations. SFVs from different categories still exhibit relatively high values, greater than 0.90, suggesting that the representation space is globally coherent while encoding only subtle inter-category variations. This block-like similarity structure closely resembles the representational similarity matrices obtained from single-neuron population activity recorded for the stimulus set used in [33], particularly in the amygdala [34]. These findings suggest that the synthetic inputs preserve a categorical organization that closely mirrors the representational structure encoded by neurons in the human medial temporal lobe.

**Figure 1:**
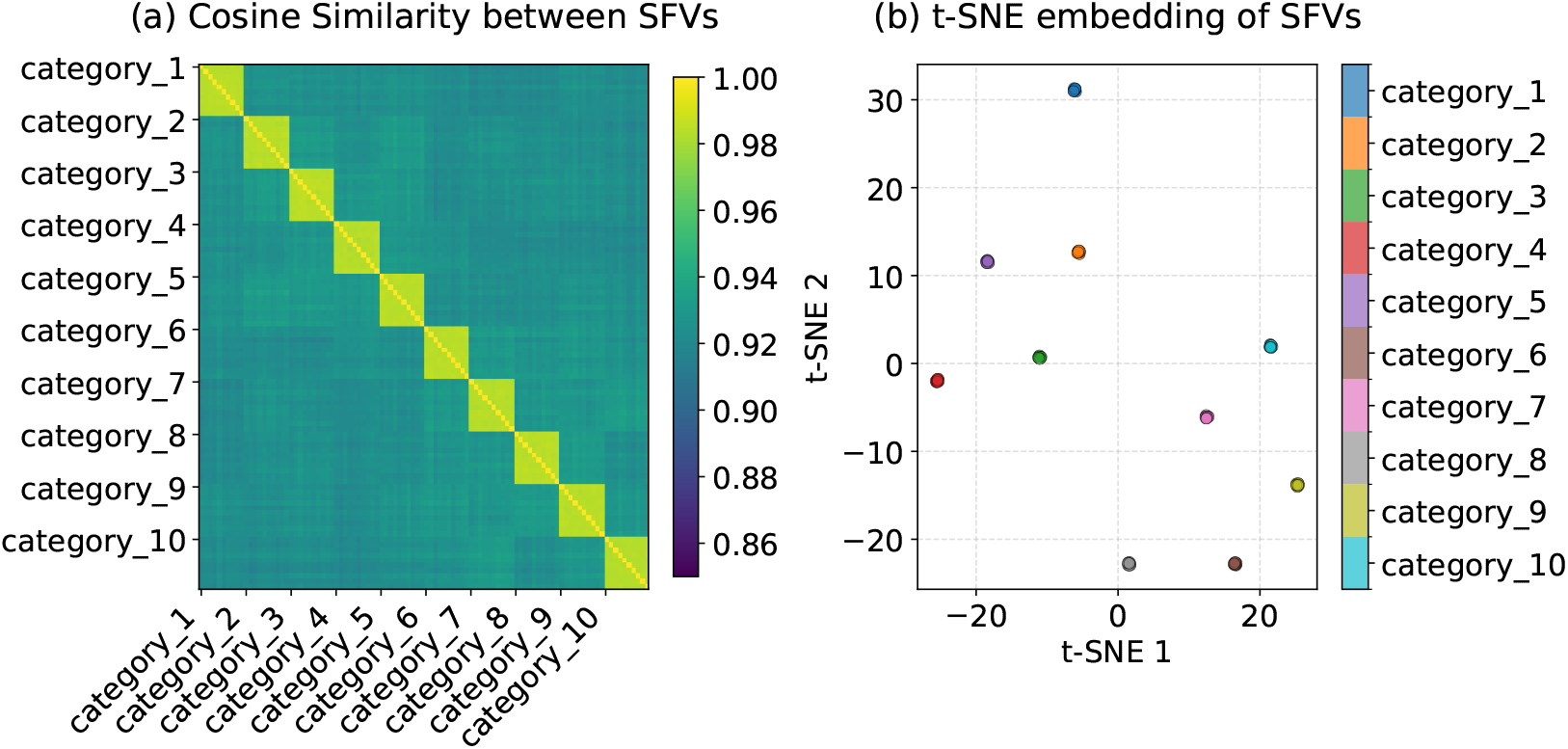
Analysis of SFV similarity and structure by category. (a) Pairwise cosine similarity among the SFVs, grouped by category. (b) tSNE projection of the SFVs into two dimensions, colored by category, revealing the clustering structure across semantic categories.

To further investigate separability, we apply *t*-distributed stochastic neighbor embedding (*t*-SNE) [41], a nonlinear dimensionality reduction technique that projects high-dimensional data into two dimensions while preserving local neighborhood structure, see the Methods section for the full formulation.

The resulting embedding (Figure 1b) reveals a well-separated structure, with samples from the same category forming compact, outlier-free clusters. Clusters corresponding to the two halves of the category set are more widely separated from each other than within each half, reflecting the presence of two higher-level meta-categories. This organization is consistent with the global structure of the embedding space, confirming that the synthetic input captures two meta-categories, each comprising five categories.

### 2.2 Population-level firing rate dynamics

A population-level analysis reveals marked differences in the temporal evolution of firing rates across the two conditions (Figure 2). In the control condition, stimulus onset elicits a sharp transient decrease in mean firing rate, lasting on average 9.3 ms (Table 1). This is followed by a rapid increase that remains elevated relative to the primed condition for approximately 49.3 ms on average (Table 2). Although both conditions exhibit a comparable onset latency, the primed condition shows an earlier return to baseline activity following the post-onset transient, indicating a shorter overall duration of the evoked response at the population level. Detailed firing-rate time courses are provided in the *Table_SourceData_SFVs*.*xlsx* file, available from the repository linked in Code availability.

**Figure 2:**
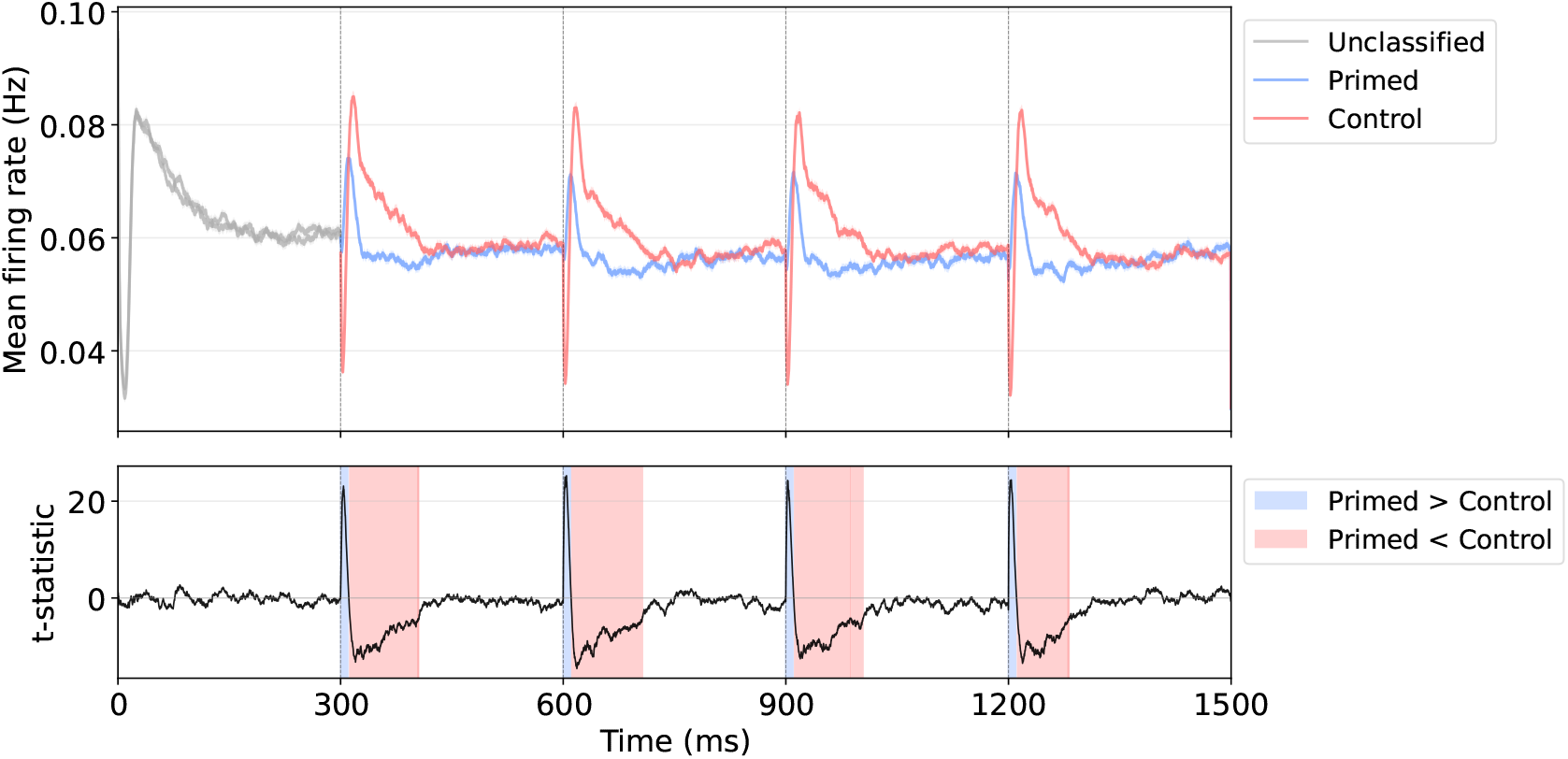
Temporal evolution of network activity. Top: Mean firing rate (in Hz) for the primed (blue) and control (red) conditions. Gray traces show the activity evoked by the first stimulus of each sequence. At this stage, the primed/control distinction has not yet emerged, and therefore these responses are not associated with either condition. The trials were randomly partitioned into two equal-sized groups solely for display purposes, highlighting the absence of a condition-specific effect before the classification becomes defined. Solid lines denote the average activity across trials. Shaded areas indicate the standard error of the mean (SEM). They are almost invisible, indicating a very small SEM. Vertical dashed lines mark stimulus onsets. Bottom: Temporal evolution of the *t*-statistic comparing the mean firing rate distributions between conditions. Blue and red bars denote time windows in which activity is significantly higher in the primed and control conditions, respectively, according to a two-sided cluster-based permutation test [24].

**Table 1:** Significant clusters in which the mean firing rate in the primed condition exceeds that in the control condition [24]. Clusters are identified using a non-parametric sign-flip permutation cluster test (*N*_perm_ = 100,000 permutations, significance threshold = 0.05). For each presentation, firing-rate activity is averaged separately in the primed and control conditions. At each sampled time point of the analyzed time series (Δ*t* = ms), a paired *t*-statistic is computed across presentations from the primed–control firing-rate differences. A cluster-defining threshold is obtained from the 95th percentile of the permutation distribution of the maximum absolute *t*-statistic. Contiguous supra-threshold time points are grouped into clusters, whose mass is defined as the sum of the absolute *t*-values across all constituent time points. Statistical significance is assessed against the null distribution of the maximum cluster mass obtained in each permutation. For each observed cluster, the *p*-value corresponds to the proportion of permutations in which the maximum cluster mass equals or exceeds the observed cluster mass, thereby controlling for multiple comparisons across time. Each row reports one significant cluster. Cluster size indicates the number of consecutive sampled time points within the cluster, whereas the reported sum of *t*-values corresponds to the signed sum of *t*-statistics across all time points belonging to the cluster. For all reported clusters, no permutation reached or exceeded the observed cluster mass; therefore, *p*-values are reported as *<* 10^*−*5^, corresponding to the resolution limit of the test, 1*/*(*N*_perm_ + 1), rather than as exactly zero.

| Time window (ms) | cluster size | sum of $t$-values | $p$-value |
| --- | --- | --- | --- |
| [300.4, 309.4] | 91 | 1442.4 | $< 10^{-5}$ |
| [600.1, 609.5] | 95 | 1672.9 | $< 10^{-5}$ |
| [900.0, 909.3] | 94 | 1467.8 | $< 10^{-5}$ |
| [1200.0, 1209.5] | 96 | 1628.0 | $< 10^{-5}$ |

**Table 2:**
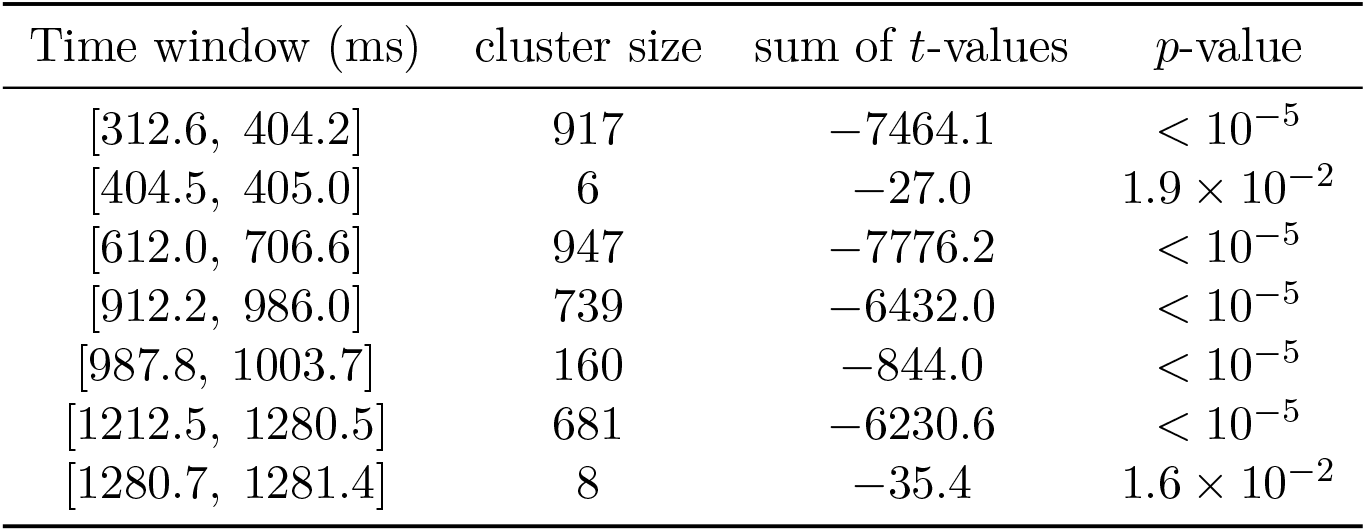
Significant clusters in which mean firing rate in the control condition exceeded that in the primed condition. Analysis parameters are as described in Table 1.

### 2.3 Neuronal recruitment across scales: from population dynamics to single-neuron onset latencies

To characterize the temporal dynamics of neuronal recruitment, we quantify the latency required for a predefined fraction of the population to reach the spiking threshold. We select activation levels of 10% and 15% to sample distinct stages of population recruitment. The 10% threshold reflects an early activation phase, capturing the fastest-responding neurons driven predominantly by strong feedforward input. The 15% threshold, in contrast, represents a later phase of recruitment, involving a broader set of neurons.

At 10% activation, neurons in the primed condition exhibit significantly shorter latencies than those in the control condition (Figure 3a, Table 3). At 15% activation, however, this latency advantage reverses, with the control condition reaching the threshold faster than the primed one (Figure 3b, Table 4). This reversal suggests that the early and late recruitment phases are governed by distinct dynamics, which we examine further in the Discussion.

**Figure 3:**
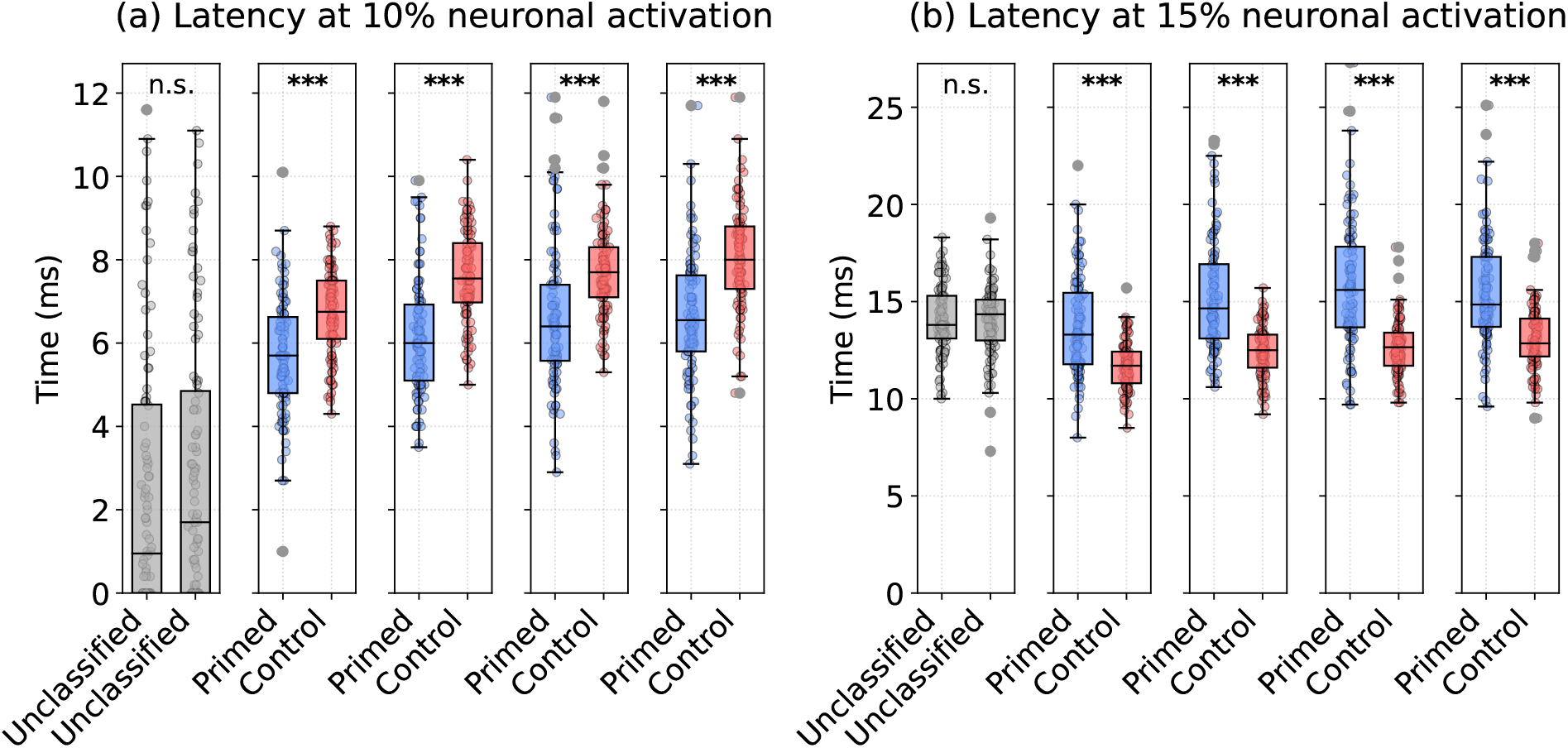
Neuronal recruitment dynamics. Box plots show the distribution of activation latencies, defined as the first time at which a predefined percentage of neurons reaches the spiking threshold, for (a) a 10% and (b) a 15% threshold. For the first time window (gray), conditions are not classified. For subsequent windows, primed and control conditions are shown in blue and red, respectively. The central line of each box indicates the median, box edges represent the 25th and 75th percentiles (interquartile range, IQR), whiskers extend to the most extreme values within 1.5 *×* IQR. Individual data points are shown as semi-transparent dots. Statistical significance between primed and control conditions is assessed using the Mann–Whitney *U* -test [29] with Holm correction [2] for multiple comparisons across time windows. Asterisks indicate significance levels: *p <* 0.001 (***) and not significant (n.s.).

**Table 3:** Population activation latency at the 10% threshold. Median latency (ms) and interquartile range [Q1, Q3] are reported for the primed and control conditions in each time window. Statistical comparisons use the Mann–Whitney *U* -test [29] with Holm correction [2] (significance threshold = 0.05). The reported *p*-value corresponds to the probability of observing a difference at least as extreme as measured, under the null hypothesis that latencies in the two conditions are drawn from the same distribution.

| Time window (ms) | primed | control | $U$ | $p$-value |
| --- | --- | --- | --- | --- |
|  | median [Q1,Q3] | median [Q1,Q3] |  |  |
| [300, 600] | 5.7 [4.8, 6.6] | 6.8 [6.1, 7.5] | 2707.0 | $4.2 \times 10^{-8}$ |
| [600, 900] | 6.0 [5.1, 6.9] | 7.6 [7.0, 8.4] | 2040.5 | $2.4 \times 10^{-12}$ |
| [900, 1200] | 6.4 [5.6, 7.4] | 7.7 [7.1, 8.3] | 2640.5 | $2.4 \times 10^{-8}$ |
| [1200, 1500] | 6.6 [5.8, 7.6] | 8.0 [7.3, 8.8] | 2285.5 | $1.3 \times 10^{-10}$ |

**Table 4:** Neuronal activation latency at the 15% threshold across time windows. Reporting format and statistical procedure are as described in Table 3.

| Time window (ms) | primed | control | $U$ | $p$-value |
| --- | --- | --- | --- | --- |
|  | median [Q1, Q3] | median [Q1, Q3] |  |  |
| [300, 600] | 13.3 [11.8, 15.5] | 11.7 [10.8, 12.4] | 7501.0 | $2.0 \times 10^{-9}$ |
| [600, 900] | 14.7 [13.1, 16.9] | 12.5 [11.6, 13.3] | 8115.5 | $1.3 \times 10^{-13}$ |
| [900, 1200] | 15.6 [13.7, 17.8] | 12.7 [11.7, 13.4] | 8117.5 | $1.3 \times 10^{-13}$ |
| [1200, 1500] | 14.9 [13.7, 17.3] | 12.9 [12.2, 14.1] | 7813.5 | $1.9 \times 10^{-11}$ |

The population-level analysis captures when a critical fraction of neurons becomes active but does not determine whether the experimental conditions alter the response timing of individual neurons. To address this, we perform a single-neuron analysis of onset latency, defined as the time of the first spike emitted by each neuron within a given analysis window following stimulus presentation. To ensure that detected onsets reflect newly emerging activity rather than the continuation of an ongoing response, onset latencies are computed only for presentations in which the reference interval immediately preceding the analysis window is completely silent. Here, the reference interval is defined as the final portion of the preceding temporal window, with duration equal to the onset-analysis reference window. Presentations violating this criterion are excluded from the onset-latency calculation. Only neurons exhibiting valid onset estimates in at least three SFV presentations are retained, ensuring that onset-latency estimates are derived from sufficiently sampled response profiles and reducing biases associated with sporadic firing activity. From the second time window onward, once the priming manipulation takes effect, neurons in the primed condition exhibit significantly shorter onset latencies than those in the control condition (Figure 4, Table 5), indicating a sustained reduction in single-neuron response latency under primed conditions beyond the initial stimulus-evoked transient.

**Figure 4:**
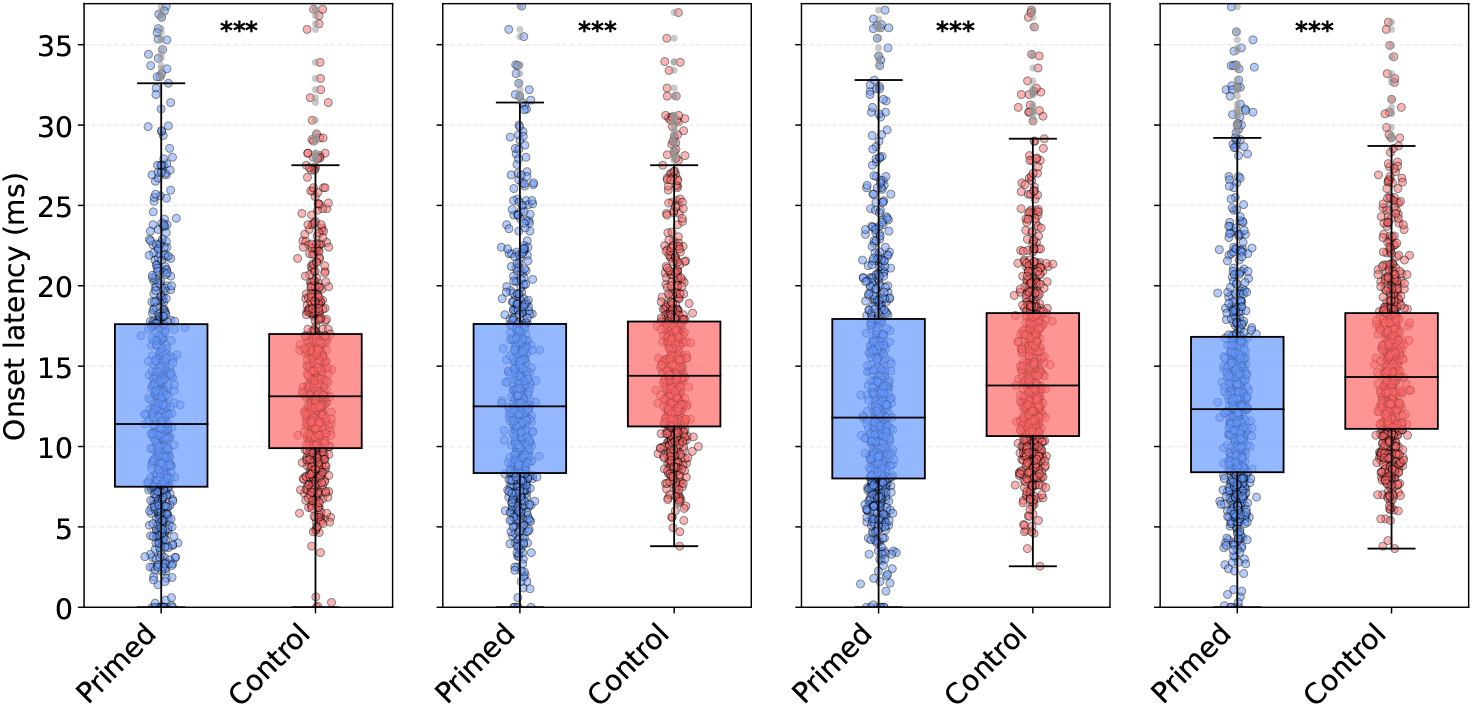
Single-neuron onset latency. Box plots show the distribution of single-neuron onset latencies under primed (blue) and control (red) conditions across time windows. In the first time window (gray), conditions are not yet differentiated. The data are reported using the same format as in Figure 3. Statistical significance is assessed using a two-tailed paired permutation test (100,000 permutations), with Holm correction for multiple comparisons across time windows (significance threshold = 0.05). Asterisks indicate significance levels: *p <* 0.001 (***) and not significant (n.s.).

**Table 5:**
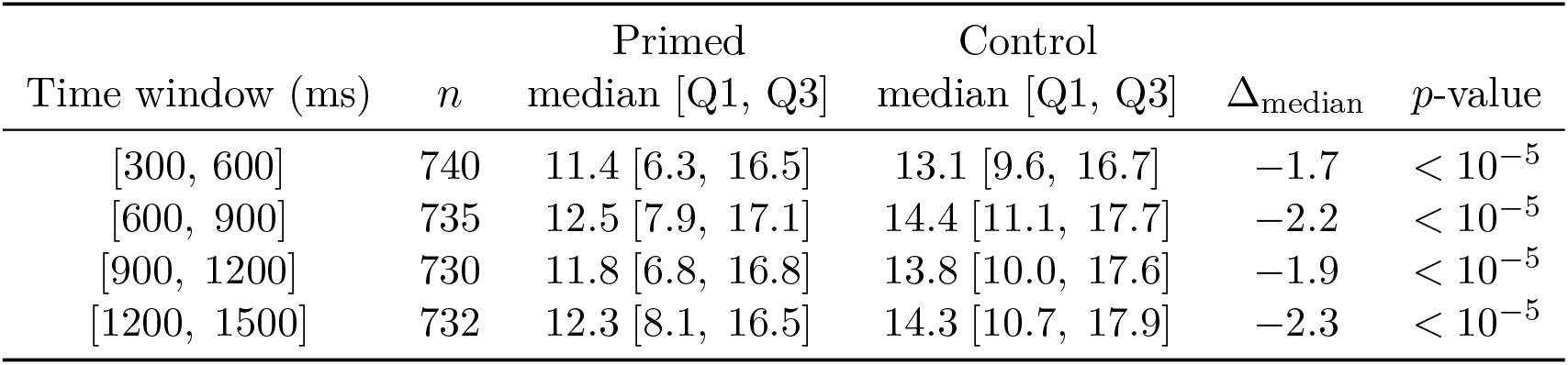
Single-neuron onset latency. For each time window: the number of included neurons (*n*, inclusion criterion: *≥* 3 active presentations under both conditions); median onset latency [Q1, Q3] for primed and control conditions; and the median within-neuron difference Δ_median_ = median(onset_primed_ *−* onset_control_). Statistical significance is assessed with a two-tailed paired permutation test (100,000 permutations): at each permutation, condition labels are exchanged within neuron pairs and the median difference is recomputed. The *p*-value is the proportion of permutations yielding an absolute median difference *≥* the observed value. All *p*-values are Holm-corrected for multiple comparisons. For all reported clusters, no permutation reached or exceeded the observed cluster mass; therefore, *p*-values are reported as *<* 10^*−*5^, corresponding to the resolution limit of the test, 1*/*(*N*_perm_ + 1), rather than as exactly zero.

| Time window (ms) | $n$ | Primed | Control | $\Delta_{\text{median}}$ | $p$-value |
| --- | --- | --- | --- | --- | --- |
|  |  | median [Q1, Q3] | median [Q1, Q3] |  |  |
| [300, 600] | 740 | 11.4 [6.3, 16.5] | 13.1 [9.6, 16.7] | -1.7 | $< 10^{-5}$ |
| [600, 900] | 735 | 12.5 [7.9, 17.1] | 14.4 [11.1, 17.7] | -2.2 | $< 10^{-5}$ |
| [900, 1200] | 730 | 11.8 [6.8, 16.8] | 13.8 [10.0, 17.6] | -1.9 | $< 10^{-5}$ |
| [1200, 1500] | 732 | 12.3 [8.1, 16.5] | 14.3 [10.7, 17.9] | -2.3 | $< 10^{-5}$ |

### 2.4 Emergence of distinct tuning phenotypes

At the single-unit level, we restrict the analysis to neurons that pass the responsiveness screening described in the Methods section. Within this subset, we observe a broad spectrum of stimulus-specific responses, with individual neurons exhibiting effects ranging from strong activation to marked suppression relative to reference activity. To quantify these deviations, we compute a stimulus-specific *Z*-score defined as

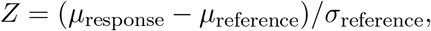

where *µ*_response_ is the mean firing rate during the 300 ms presentation window of the current stimulus, whereas *µ*_reference_ and *σ*_reference_ are the mean and standard deviation of the firing rate during the immediately preceding 300 ms stimulus-presentation window. Because stimuli are presented consecutively without inter-stimulus intervals, this preceding window is not a stimulus-free baseline but corresponds to the presentation of the immediately preceding stimulus. Accordingly, responsiveness screening and *Z*-score computation are performed only for the second through fifth stimuli of each sequence, with the first stimulus providing the context required to classify the subsequent stimulus as primed or control. Network activity reaches a stable regime approximately 200 ms after stimulus onset. Control analyses using only the final 100 ms of the preceding window, as well as analyses omitting the initial Wilcoxon screening (see Wilcoxon signed-rank test), yield unchanged results; therefore, the full preceding 300 ms window is retained as the reference interval. The underlying spiking activity is simulated with a temporal resolution of 0.1 ms.

To further characterize these response profiles, we restrict the analysis to the subset of stimuli that drive each neuron most strongly. For each neuron, stimuli are ranked in descending order of their *Z*-score under the control condition, and only the strongest are retained. This focused analysis reveals that neuronal responses segregate into two distinct functional phenotypes. The first phenotype corresponds to a sharpening-like modulation (Figure 5a), priming induces a pronounced divergence between conditions across stimulus ranks, with the primed condition exhibiting a systematically lower and more steeply modulated profile compared to control. In contrast, a second class of neurons exhibits a fatiguing-like phenotype (Figure 5b), the *Z*_primed_ and *Z*_control_ traces remain aligned across stimulus ranks, with only small divergence between conditions, as further quantified in Table 6.

**Figure 5:**
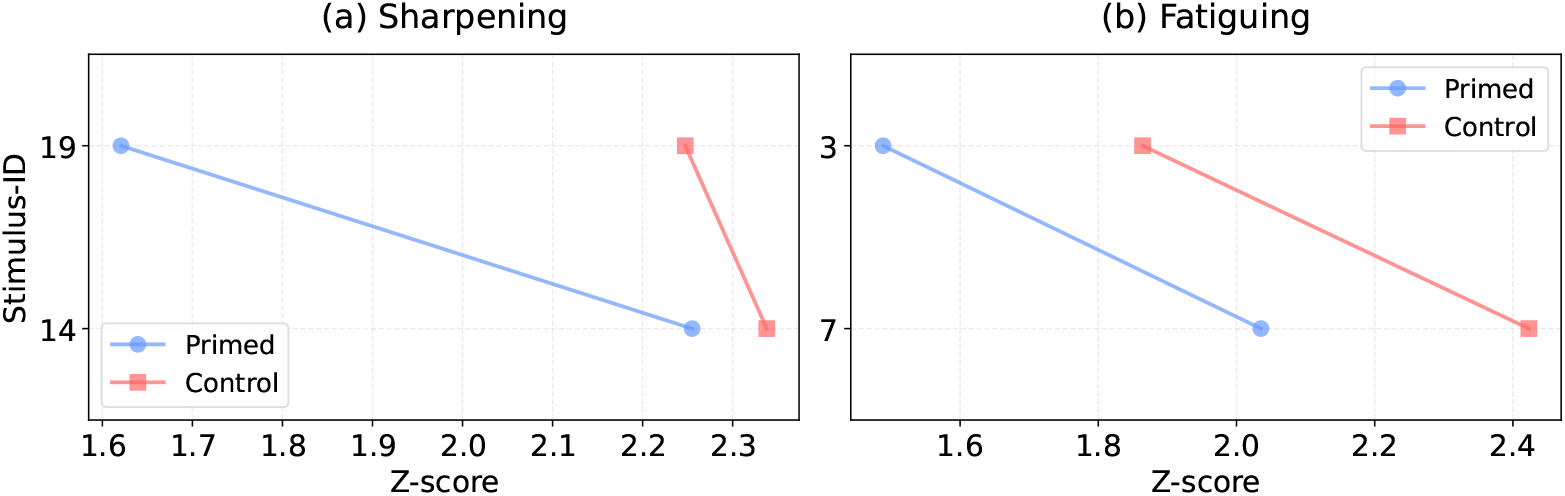
Sharpening and fatiguing response phenotypes at the single-neuron level. *Z*-scored response amplitudes for two exemplar neurons under primed (blue) and control (red) conditions. For each neuron, stimuli are ranked in descending order of their control-condition *Z*-score, from most to least effective. (a) Neuron displaying a sharpening-like phenotype, with consistently reduced and more sharply modulated responses under priming relative to control. (b) Neuron displaying a fatiguing-like phenotype, where primed and control responses remain closely aligned across stimulus ranks. Corresponding *Z*-difference (= *Z*_control_ *− Z*_primed_) values are reported in Table 6.

**Table 6:**
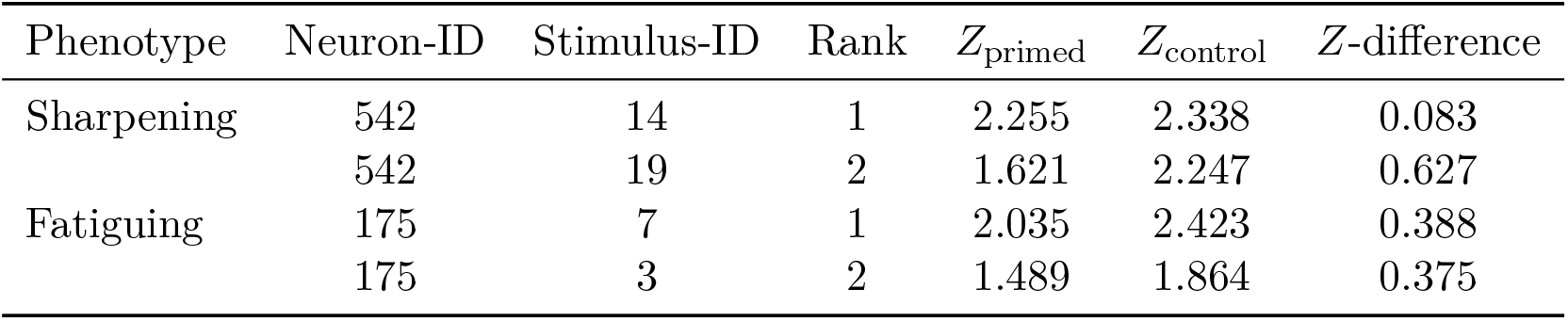
Stimulus-wise *Z*-scored responses for the representative neurons shown in Figure 5. Stimuli are ranked in descending order of the control condition *Z*-score. Reported values correspond to responses under primed and control conditions, together with the associated *Z*-difference (= *Z*_control_ *− Z*_primed_).

### 2.5 Neural response classification

To automatically classify the response behavior of individual neurons across experimental conditions, we introduce a scalar summary statistic, which we term the *delta metric* (Δ). For each neuron, the two stimuli eliciting the strongest responses are identified and ranked by decreasing *Z*-score magnitude under the control condition. The delta metric is then defined as

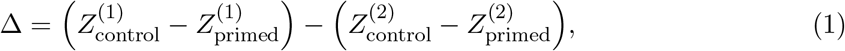

where superscripts (1) and (2) denote the first- and second-ranked stimuli, respectively. A negative Δ indicates that the leading stimulus becomes relatively more selective under priming (sharpening), whereas a value near zero indicates a symmetric attenuation of responses across the two top stimuli (fatiguing).

Prior to computing Δ, neurons are retained only if they pass a two-stage selectivity criterion. In the first stage, a neuron must exhibit |*Z*| *>* 2 in both conditions for at least one stimulus. In the second stage, the same neuron must show |*Z*| *>* 1 in both conditions for at least two distinct stimuli. This joint criterion ensures that only genuinely stimulus-selective neurons enter the classification pipeline.

Classification boundaries are defined using the median absolute deviation (MAD), a robust dispersion measure unaffected by outliers or heavy distributional tails, i. e.,

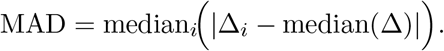

The classification tolerance is set to *τ* = 0.5 *×* MAD, and individual neurons are assigned to one of two functional categories: sharpening if Δ *< −τ*, while fatiguing if |Δ| *≤ τ* . Values of Δ *> τ* are not assigned to either response phenotype. Such values correspond to a qualitatively distinct response regime in which a neuron’s preferred stimulus is suppressed more strongly than a less-preferred stimulus. As this response pattern is not typically reported in experimental studies of repetition suppression, neurons with Δ *> τ* are excluded from subsequent analyses.

A network-level label is then assigned by majority vote: a configuration is designated *sharpening* if the count of sharpening neurons exceeds that of fatiguing neurons, and *fatiguing* otherwise. Configurations in which the majority of classified neurons have Δ *> τ*, or in which sharpening and fatiguing counts are exactly equal (ties), are discarded from comparative analyses. The network driven by the reference SFVs (see Synthetic feature vectors) is classified as sharpening.

### 2.6 From single-neuron dynamics to network-level behavior across a systematic input sweep

To assess whether and how the structure of the input patterns shapes the collective behavior of the network, we perform a systematic sweep over 200 input triplets. Each triplet is defined by three parameters (*α, β, ε*), which jointly determine the statistical structure of the SFVs (see Synthetic feature vectors). Across all 200 conditions, the ordering of the SFVs presented to the network remains fixed, only the internal vector structure varies as a function of the three parameters.

Each simulated network is classified as sharpening or fatiguing using the majority-vote criterion described above. Configurations for which the majority of classified neurons have Δ *> τ*, and configurations resulting in exact ties, are excluded. This procedure yields 42 sharpening and 29 fatiguing configurations retained for all subsequent comparisons, whereas 85 configurations do not reach a majority in either category and 34 result in a tie, both excluded from further analyses. The resulting phenotype assignments are provided in the *Phenotypes*.*xlsx* file, available from the repository linked in Code availability.

The distribution of labels across the parameter space reveals a clear dependence of network behavior on input structure (Figure 6a). Statistical comparison of the parameter values associated with each network class reveals significant differences for *α* and *β*, but not for *ε* (Figure 6c, Table 7). Specifically, networks classified as sharpening operate under significantly higher values of both *α* and *β* compared to those classified as fatiguing, while the two groups are indistinguishable with respect to *ε*.

**Figure 6:**
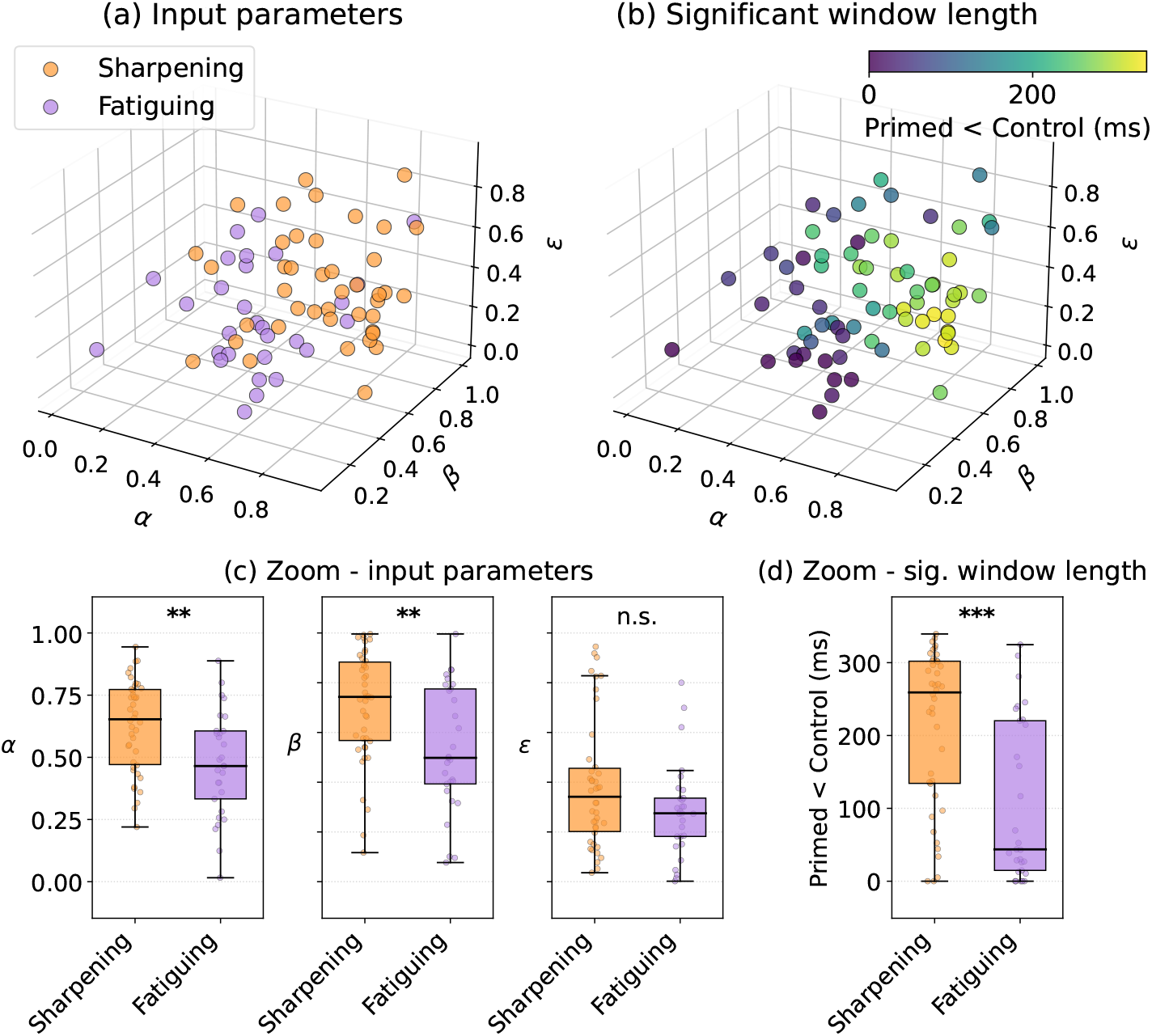
Parameter-dependent classification of network behavior. (a) Three-dimensional scatter plot of the retained simulated configurations in the (*α, β, ε*) parameter space, color-coded by majority-vote network label (orange: sharpening, *n* = 42; purple: fatiguing, *n* = 29). (b) Same configurations color-coded by the duration (ms) of the window in which the mean firing rate of the primed condition is statistically significantly below the control condition one. (c) Box plots of the three input parameters stratified by network class. (d) Box plots of the suppression window duration stratified by network class. In all box plots, the central line denotes the median, box edges denote the first and third quartiles, and whiskers extend to 1.5 *×* IQR, individual data points are overlaid with jitter. Asterisks denote significance levels (Mann–Whitney *U* -test, two-sided), *p <* 0.001 (***), *p <* 0.01 (**), and not significant (n.s.).

**Table 7:** Statistical comparison of input parameters between sharpening (*n* = 42) and fatiguing networks (*n* = 29). Results of Mann–Whitney *U* -tests (two-sided) comparing the distributions of *α, β*, and *ε* between the two network classes. Values are reported as median [Q1, Q3].

| Parameter | sharpening | fatiguing | $U$ | $p$-value |
| --- | --- | --- | --- | --- |
|  | median [Q1, Q3] | median [Q1, Q3] |  |  |
| $\alpha$ | 0.653 [0.471, 0.773] | 0.465 [0.333, 0.606] | 861.0 | $3.3 \times 10^{-3}$ |
| $\beta$ | 0.743 [0.568, 0.883] | 0.498 [0.393, 0.775] | 857.5 | $3.7 \times 10^{-3}$ |
| $\epsilon$ | 0.342 [0.202, 0.456] | 0.275 [0.182, 0.336] | 740.5 | $1.2 \times 10^{-1}$ |

To further characterize the relationship between network classification and the temporal dynamics of the priming effect, we examine the duration of the time window in which the primed condition is statistically significantly below the control condition, i. e., the window of repetition suppression, as described in Section 2.2. Sharpening networks exhibit a significantly longer suppression window compared to fatiguing networks (Mann–Whitney *U* -test, *U* = 928.0, *p* = 1.93 *×* 10^*−*4^), with median durations [Q1, Q3] of 259.2 [134.3, 302.0] ms and 43.7 [14.8, 220.3] ms, respectively (Figure 6b and d).

Finally, to evaluate whether the model as a whole captures sharpening behavior at the population level, we examine the distribution of Δ values pooled across all valid neurons and all conditions. The distribution is significantly non-normal (Shapiro–Wilk test, *p* = 0.013), and the negative median, Δ_median_ = *−*0.125, indicates a systematic bias towards sharpening across the full parameter space.

## 3 Discussion

We developed a biologically grounded spiking network model of visual semantic adaptation and use it to investigate the microcircuit mechanisms underlying two distinct neural adaptation phenotypes: sharpening and fatiguing. Our results show that both phenotypes emerge naturally from recurrent dynamics shaped by triplet-STDP, and that the balance between them is governed by the statistical structure of the sensory input rather than by qualitative differences in network architecture.

At the population level, priming consistently reduces mean firing rates relative to the control condition across all four stimulus presentation windows (Figure 2, Table 2). This reduction is preceded by a brief transient in which the control condition temporarily drops below the primed condition, reflecting the early post-onset dynamics of the network response. The sustained suppression that follows is consistent with the interpretation that representations learned through triplet-STDP are reactivated more efficiently when preceded by a related stimulus: reactivation occurs in both the primed and control conditions, but proceeds faster and at lower energetic cost when the preceding stimulus shares semantic structure with the current one [4, 17]. The temporal profile of this suppression, characterized by an earlier onset and a reduced peak amplitude in the primed relative to the control condition, closely mirrors the event-related potential compression reported in iEEG [33], suggesting that the model captures the macroscopic signature of adaptation through the aggregate dynamics of its spiking populations.

The reversal of the latency advantage between the 10% and 15% activation thresholds (Figure 3) provides insight into the structure of the priming effect at the subpopulation level. At early recruitment stages, primed stimuli activate a small cohort of highly selective neurons faster than control stimuli, consistent with a facilitation of the initial feedforward sweep through recurrent connections strengthened during learning [25]. At later stages, the broader recruitment of less selective neurons is delayed under priming. This delay reflects adaptation-driven suppression mediated by spike-frequency adaptation and short-term synaptic depression, together with lateral inhibitory interactions. The rapid engagement of highly selective neurons under priming suppresses neighboring, less selective units through recurrent inhibitory connections, thereby reducing their excitability and postponing their recruitment relative to the control condition.

This two-stage pattern is entirely consistent with the sharpening hypothesis, in which priming selectively amplifies high-selectivity responses while attenuating those of lower specificity [27, 42], and has been previously described in computational models of competitive neural coding [27]. A qualitatively similar pattern is observed in the DINO embedding condition (see the DINO feature embeddings appendix), suggesting that the mechanism is intrinsic to the recurrent circuit rather than dependent on the specific input statistics.

The single-unit analysis reveals that the two adaptation phenotypes co-exist within the same simulated network: individual neurons exhibiting a sharpening-like divergence between primed and control tuning curves varies between neurons showing a fatiguing-like parallel attenuation (Figure 5). This heterogeneity mirrors findings in the human medial temporal lobe, where sharpening is predominantly reported in the amygdala and fatiguing in the hippocampus and parahippocampal cortex [33], suggesting that circuit-level differences in recurrent connectivity and adaptation dynamics, rather than fundamentally different neural mechanisms, may account for regional specificity. In our model, the balance between phenotypes within a given network is captured by the majority-vote label assigned to each configuration: the signed Δ metric has a negative median across the full parameter sweep, indicating that sharpening is the dominant mode across the explored input space. This result is quantitatively consistent with the response profiles reported in single-neuron recordings [33], where sharpening-compatible patterns are particularly prevalent in the amygdala.

The systematic parameter sweep over 200 input triplets reveals that the expression of sharpening versus fatiguing is not fixed but depends sensitively on input structure (Figure 6, Table 7). Specifically, higher values of *α* (intra-category similarity) and *β* (inter-category dissimilarity) favor sharpening, whereas the contraction parameter *ε* does not significantly discriminate between phenotypes. This pattern is interpretable in terms of the representational geometry of the input: high *α* produces tightly clustered category representations, increasing within-category similarity; high *β* increases the distance between cluster centroids, making category boundaries more salient. Under these conditions, the recurrent network is confronted with a highly structured input space in which selective strengthening of the connections encoding the most discriminative stimulus features becomes both possible and advantageous. The resulting connectivity, shaped by triplet-STDP, preferentially enhances responses to the most effective stimuli while suppressing responses to less preferred ones, consistent with sharpening. Conversely, when *α* and *β* are low, the representational geometry is diffuse, and the network defaults to a more uniform adaptation, consistent with fatiguing. These findings suggest a direct link between representational geometry and functional adaptation mode, extending earlier theoretical proposals that stimulus discriminability governs the transition between sharpening and fatiguing [19, 27]. It is worth noting that the majority-vote classification scheme retains a subset, 71 out of 200 (35.5%), of the simulated configurations for direct comparison between sharpening and fatiguing networks, while the remaining ones do not achieve a majority classification in either class. This exclusion is a direct consequence of the binary classification framework adopted here, which is designed specifically to test the hypothesis that input geometry governs the sharpening–fatiguing continuum, rather than to exhaustively characterize every possible adaptation regime. Notably, a portion of these unclassified configurations is dominated by neurons with Δ *> τ*, the regime in which a neuron’s preferred stimulus is suppressed more strongly than a less-preferred one, raising the possibility that the same network architecture may also support enhancement-like dynamics.

The weaker adaptation effects observed with DINO embeddings compared to synthetic inputs (see the DINO feature embeddings appendix) can be understood in terms of the same geometric framework. As shown in Figure 7 and Figure 8, DINO features form a more compact and less segregated representational space, with smaller inter-category distances and weaker cluster structure. Under these conditions, the network operates in a regime of low effective *α* and *β*, precisely the regime in which our parameter sweep predicts weaker sharpening and reduced suppression windows. This convergence between the geometric analysis and the empirical observations validates the synthetic input framework as a useful tool for systematically exploring the conditions that promote adaptation in recurrent networks. An important direction for future work is to improve the alignment between synthetic and natural image representations, for example by incorporating semantic structure from large-scale language models.

**Figure 7:**
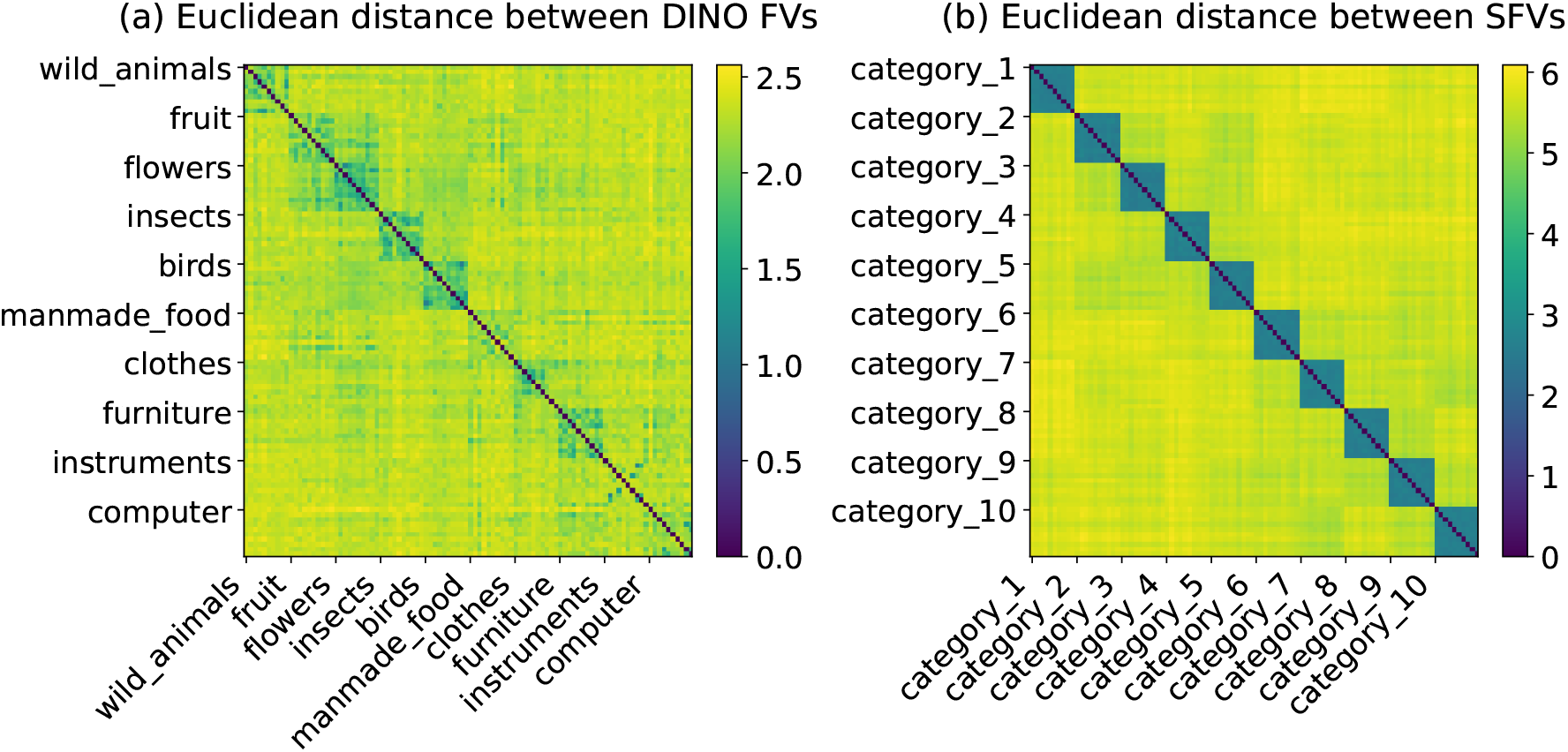
Comparison of input feature representations. Heatmaps show the pairwise Euclidean distances among DINO FVs (a) and SFVs (b). Vectors are grouped by category, and only the label of the first occurrence of each stimulus is displayed.

**Figure 8:**
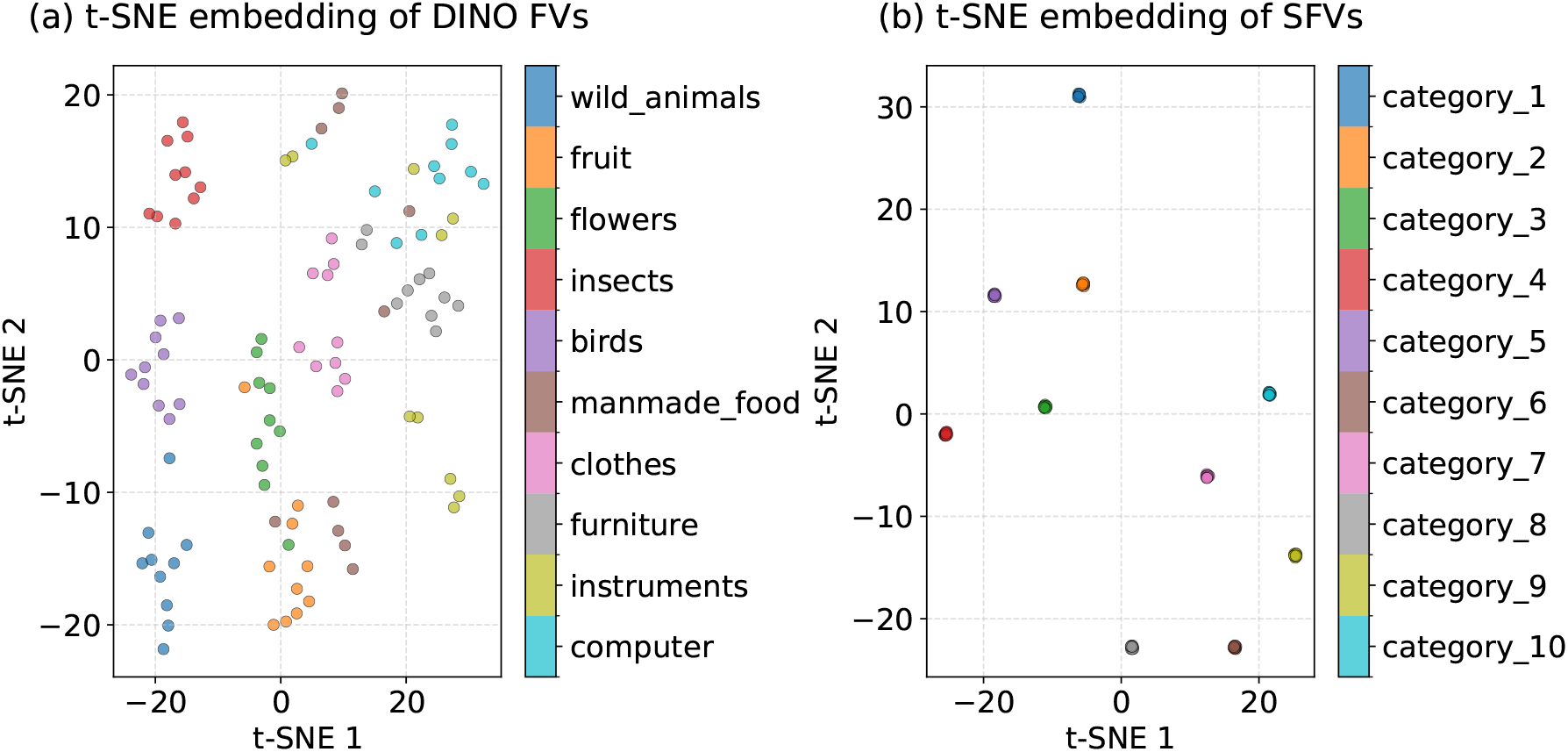
Comparison of input feature representations. *t*-SNE embeddings of DINO FVs (a) and SFVs (b) projected onto two dimensions. Each color represents a specific category.

Our results are broadly consistent with predictive coding frameworks in which repeated stimuli generate weaker prediction errors, leading to suppressed neural responses [15]. In this interpretation, learning through triplet-STDP establishes a prior over stimulus sequences, such that the primed condition produces a smaller mismatch between the network’s expectations and the incoming stimulus. Sharpening would then reflect a more precise prior that concentrates prediction onto the most diagnostically relevant stimulus features, whereas fatiguing would reflect a less specific prior that uniformly attenuates responses. Our finding that input geometry governs this transition is consistent with the proposal that prior precision is itself a function of representational discriminability [15, 27].

Several aspects of the present framework provide opportunities for future extensions. First, the current network architecture does not explicitly incorporate anatomically differentiated subregions, and therefore does not aim to reproduce the full regional specificity of adaptation phenotypes reported across the amygdala, hippocampus, and parahippocampal cortex [33]. Second, the triplet-STDP rule adopted here provides a phenomenological description of synaptic plasticity, without explicitly modeling mechanisms such as presynaptic short-term depression or neuromodulatory modulation, both of which have been implicated in fatiguing-like adaptation [13, 32]. Integrating these processes could further refine the mechanistic interpretation of the observed dynamics. Third, the unclassified configurations dominated by Δ *> τ* neurons point to a possible enhancement-like regime that the present binary classification was not designed to capture, and which merits dedicated investigation. Fourth, the model generates experimentally testable predictions regarding the influence of stimulus statistics on adaptation dynamics. Future single-unit recordings in humans could systematically manipulate the *α* and *β* parameters governing stimulus similarity and meta-category structure, providing a direct test of whether the predicted changes in priming-related neural responses are observed in biological neural populations.

Collectively, our results demonstrate that response sharpening and fatiguing are not discrete mechanisms requiring fundamentally different circuit architectures, but rather emergent modes of a single recurrent spiking network whose expression is determined by the geometry of the sensory input. This provides a unifying mechanistic account of adaptation diversity across cortical regions and stimulus conditions, and offers a principled computational framework for generating and testing experimental predictions about the neural basis of visual semantic priming.

## 4 Methods

### 4.1 Network design

Let *N* denote the total number of neurons in the network. The population is divided into an excitatory group of size 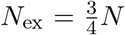 and an inhibitory group of size 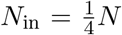. The network comprises four types of synaptic connections: excitatory-to-excitatory (ex→ex), excitatory-to-inhibitory (ex *→* in), inhibitory-to-excitatory (in *→* ex), and inhibitory-to-inhibitory (in *→* in). These connections are randomly established according to fixed connection probabilities consistent with those reported in [14]: *p*_ex*→* ex_ = 0.065, *p*_ex*→* in_ = 0.200, *p*_in*→*ex_ = 0.275, *p*_in*→* in_ = 0.100. Synaptic weights are initialized from a uniform distribution in the range [0, 0.25] for excitatory synapses and [*−*0.25, 0] for inhibitory ones.

### 4.2 Neuronal model

The subthreshold dynamics of a neuron’s membrane potential, in response to an input current, are described by the AdEx model with conductance-based synaptic interactions. All simulations are implemented using the `Brian2` simulator [38] and executed in standalone mode, where the model is compiled into optimized code to enhance computational efficiency. Differential equations are numerically integrated using the forward Euler method with a simulation time step of 0.1 ms. For a postsynaptic neuron *i*, the membrane potential evolves according to

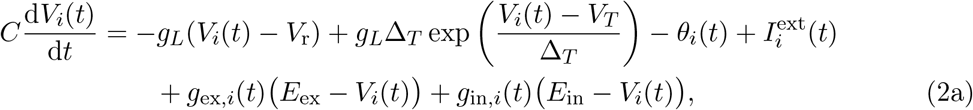

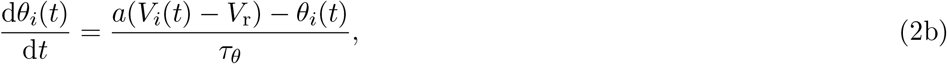

where *C* = 281 pF is the membrane capacitance, *g*_*L*_ = 30 nS the leak conductance, *V*_r_ = *−*70.6 mV the resting potential, Δ_*T*_ = 2 mV the slope factor, and *V*_*T*_ = *−*50.4 mV the threshold potential. The variable *θ*_*i*_(*t*) represents spike-frequency adaptation with time constant *τ*_*θ*_ and *a* is the subthreshold adaptation conductance, while 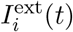 represents the external input current.

Each neuronal population is characterized by a specific set of parameters reflecting their electrophysiological profiles in accordance with [6]. For excitatory neurons, the parameters are *τ*_*θ*_ = 144 ms and *a* = 4 nS. For inhibitory neurons, we set *τ*_*θ*_ = 300 ms, and 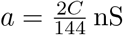.

Synaptic interactions are modeled via conductance-based inputs. The variables *g*_ex,*i*_(*t*) and *g*_in,*i*_(*t*) denote the total excitatory and inhibitory synaptic conductances received by neuron *i*, respectively. Their temporal evolution follows first-order exponential decay

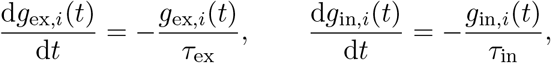

with *τ*_ex_ = 5 ms and *τ*_in_ = 10 ms. The corresponding synaptic reversal potentials are set to *E*_ex_ = 0 mV and *E*_in_ = *−*80 mV for excitatory and inhibitory synapses, respectively. The excitatory reversal potential is consistent with experimental measurements of AMPA-dominated synapses [9] and is commonly adopted in single-compartment models as a biologically reasonable approximation [30].

Synaptic transmission is not instantaneous. To account for finite axonal and dendritic conduction times, we assign a small randomly distributed delay to each synapse. The ex *→* ex and ex *→* in synapses have delays drawn uniformly in [1, 1.5] ms, whereas in *→* ex and in *→* in synapses are assigned delays in [0.5, 1] ms [5, 28].

Let *w*_*ij*_(*t*) denote the synaptic weight from presynaptic neuron *j* to postsynaptic neuron *i*. When neuron *j* emits a spike at time 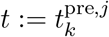, the synaptic weight produces an instantaneous increment of the postsynaptic conductance:

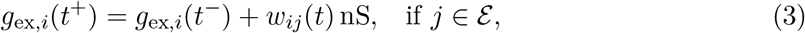

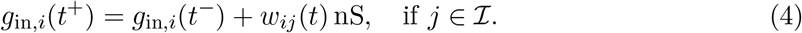

where *ℰ* and *ℐ* denote the sets of excitatory and inhibitory neurons, respectively. The synaptic weights *w*_*ij*_(*t*) modulate the amplitude of the conductance jump, while the sign of the resulting synaptic current is determined by the corresponding reversal potential.

A spike is emitted by neuron *i* when *V*_*i*_(*t*) exceeds the cutoff threshold *V*_cut_ = *V*_*T*_ + 5Δ_*T*_ . At this moment, the membrane potential is reset and the adaptation variable is increased. More precisely, we obtain jumps in the functions *V*_*i*_(·) and *θ*_*i*_(·), i. e.,

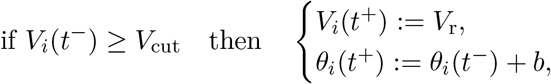

where *V*_r_ is the resting potential and *b* is the spike-triggered increment of adaptation. Depending on the type of neuron, we choose different values for *b*, namely, *b* = 0.0805 nA for excitatory neurons, and *b* = 0 nA for inhibitory neurons.

The synaptic weights during the testing phase are static, while during the learning phase, *w*_*ij*_(*t*) evolves according to a triplet-based spike timing dependent plasticity rule, driven by the precise timing difference between presynaptic and postsynaptic spikes timing. The full formulation of this mechanism is presented in the Learning phase subsection.^1^

Finally, in order to obtain a unique solution of the differential equations in (2), its initial conditions are set with the membrane potential at the resting value, *V*_*i*_(0) = *V*_r_, the adaptation variable at zero, *θ*_*i*_(0) = 0, representing a neuron at rest prior to the application of any input current, and with the synaptic conductances initialized to zero, *g*_ex,*i*_(0) = *g*_in,*i*_(0) = 0.

### 4.3 External input current

The external input current 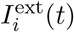 represents a neuron-specific, time-dependent drive that constitutes the principal pathway through which external signals are delivered to the network. It is first implemented as a transient oscillatory input to perturb network dynamics during initialization, and subsequently as a temporally structured sequence of high-dimensional feature vectors encoding visual (see the DINO-derived embeddings subsection) and synthetic stimuli (see the Synthetic feature vectors subsection).

Formally, the external input current is defined as

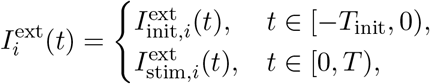

where *T*_init_ denotes the duration of the initialization phase, and *T* denotes the total duration of the temporal window assigned to each feature presentation. In the following, we describe the construction of the initialization current 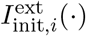 and the two alternative realizations of the stimulus-driven input 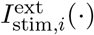 employed in the simulations.

#### Initialization input

During the initial *T*_init_ = 100 ms of each simulation, all neurons receive a sinusoidal input current with randomized phase and scaled amplitude (in nanoamperes, nA), defined as

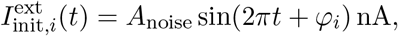

where *A*_noise_ = 0.98 sets the amplitude of the sinusoidal pre-stimulation input. Although all neurons are subject to the same baseline oscillatory component, each neuron is assigned an independent phase *ϕ*_*i*_ *∼ U*([0, 2*π*]), i. e., sampled uniformly at random between 0 and 2*π*. This randomized phase assignment induces heterogeneous transient dynamics across the network during the initialization period. To match the scale of other network inputs, which are normalized between 0 and 1, the resulting currents are clipped to this interval.

#### Stimulus-driven input

After the initialization phase, the network is evaluated under two distinct types of input. The first is derived from high-level visual features extracted from the image dataset used in [33], while the second one is generated from synthetic feature vectors defined by three key parameters, whose appropriate combination allows us to approximate the features structure derived from the former.

In this regime, the external input current is defined by

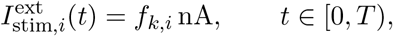

where **f**_*k*_ = [*f*_*k*,1_, …, *f*_*k,N*_]^*T*^ *∈* [0, 1]^*N*^ denotes the feature vector associated with stimulus *k*, and *N* is both the number of neurons in the network and the dimensionality of the feature vectors, establishing a one-to-one mapping between feature channels and neurons. Both the values of *T* and the temporal order in which the feature vectors **f**_*k*_ are presented depend on the experimental phase (learning or testing) and are detailed in the Learning phase subsection and Results section.

If a network with a different number of neurons is required, the feature vectors can be linearly transformed to match the network size. Specifically, when the number of neurons exceeds the feature dimensionality, each feature can be distributed across multiple neurons, whereas when the network has fewer neurons than features, the vectors can be downsampled or projected via dimensionality reduction techniques, preserving the essential structure of the input representation.

#### 4.3.1 DINO-derived embeddings

High-level visual features are extracted from the *S* = 100 images employed in the experiment reported in [33], organized into two semantic meta-categories, each comprising five categories of ten images.

Each image *i* is first resized to 224 *×* 224 pixels to match the input resolution expected by the DINO vision transformer model [8], a self-supervised framework capable of learning abstract visual representations without labeled data. The resized images are then converted into tensors **Z**_*i*_ *∈* [0, 1]^3*×*224*×*224^, where the first dimension corresponds to the three RGB color channels which are normalized channel-wise according to

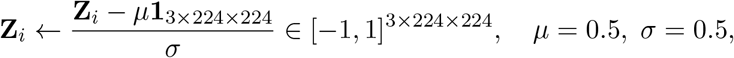

where **1**_3*×*224*×*224_ denotes the 3 *×* 224 *×* 224 tensor of all ones. The choice of *µ* and *σ* is geometric: *µ* corresponds to the midpoint of the unit interval, while *σ* scales the range so that the maximum absolute deviation from zero is 1, in accordance with the requirements of the pre-trained model used as input to DINO.

The model processes each image through a hierarchy of transformer layers that capture both local and global visual structure, yielding a high-dimensional embedding for each image. These embeddings are aggregated into a feature matrix 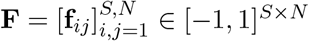, where each row encodes the representation of a single image. Its rows are ordered such that consecutive entries correspond to images from the same category, before advancing to the subsequent category within the same overarching meta-category.

In our simulation, we use *N* = 768 neurons, corresponding exactly to the dimensionality of the DINO feature vectors. To ensure compatibility with current-based network models [31], the features are normalized to the unit interval by first subtracting the minimum of the entries of **F**, i. e., **F**_min_ := min_*i,j*_ **f**_*ij*_, from all entries and dividing by the maximum of the entries, **F**_max_ := max_*i,j*_ **f**_*ij*_, so that **F** is replaced by

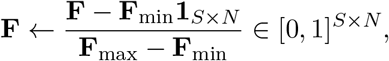

where **1**_*S×N*_ denotes the *S × N* matrix of all ones. During network simulations, stimulus-driven input currents are generated by sequentially presenting selected rows of **F**.

#### 4.3.2 Synthetic feature vectors

To simulate structured inputs analogous to the DINO embeddings described above, we generate a synthetic dataset of *S* = 100 high-dimensional vectors **f**_*i*_ *∈* [0, 1]^*N*^, with *N* = 768. The dataset is hierarchically organized into two meta-categories, each containing five categories of ten samples, resulting in a block structure that mirrors the categorical hierarchy of the DINO embeddings.

Each category *k* is associated with a reference vector (centroid) **c**_*k*_ *∈* ℝ^*N*^, generated by adding a random perturbation vector ***δ***_*k*_ to a global base vector **g** *∈* [0, 1]^*N*^ whose entries are sampled independently from a uniform distribution in [0, 1], i. e.,

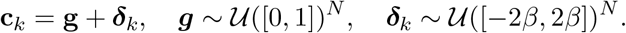

The parameter *β ∈* [0, 1] controls the spread of the category centroids around the global base, i. e., the degree of inter-category dissimilarity. The feature vectors **f**_*i*_ within each category *k* are generated as Gaussian perturbations with zero mean and covariance *σ*^2^**I**_*N*_ around **c**_*k*_, that means

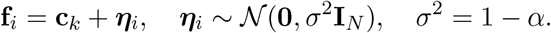

The parameter *α ∈* [0, 1] controls intra-category similarity, larger values of *α* yield lower variance and hence more tightly clustered samples.

To further impose a hierarchical separation between the two meta-categories, we partition the synthetic dataset of feature vectors **f**_*i*_ into two disjoint groups of equal size (*M* = 50 samples each), corresponding to the five categories of each meta-category. For each group, the centroid is computed as the element-wise mean

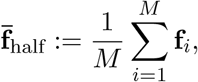

Each feature vector is then shifted towards the centroid of its respective group according to

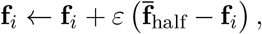

where the parameter *ε ∈* [0, 1] controls the strength of the contraction. Larger values of *ε* pull the samples more strongly toward their group centroid, thereby reducing intra-group variability and thereby enhancing the relative separability between the two meta-categories. Conversely, setting *ε* = 0 leaves the vectors unchanged.

Following the procedure described in the DINO-derived embeddings subsection, synthetic feature vectors are organized into a matrix **F** *∈* ℝ^*S×N*^, normalized to the unit interval, and use to generate the external input currents *I*^ext^ to drive the network during simulations.

##### Selection of synthetic input parameters

To identify the parameter regime that maximizes priming-induced repetition suppression, we optimize the triplet (*α, β, ε*) using differential evolution (DE) [39], a gradient-free global optimization algorithm well suited to non-convex objective functions. For each candidate triplet (*α, β, ε*) ∈ [0, 1]^3^, the full simulation pipeline is executed, including synthetic feature generation, associative-network training through triplet-STDP (see Learning phase), and evaluation on *N*_seq_ = 10 randomly generated stimulus sequences balanced across primed and control conditions. Population firing rate dynamics are compared using a cluster-based permutation test [24] (10,000 permutations, significance threshold = 0.02, minimum cluster size = 5 bins). We quantify the cumulative duration of significant temporal clusters for which firing activity in the primed condition is lower than in the control condition, denoted *C*_primed*<*control_, and optimize

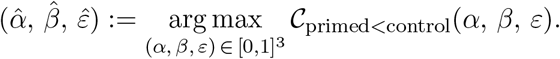

Optimization is performed using the `rand1bin` strategy (population size multiplier = 10 (30 individuals), mutation factor *F* = 0.8, crossover probability CR = 0.9, 20 generations and random seed = 42). Parameters are rounded to three decimal places prior to evaluation to ensure reproducibility. The optimization converges to

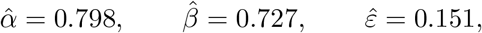

yielding a maximal *C*_primed*<*control_ value of 339.4 ms. Unless otherwise specified, these values are used in the reference simulations presented in the Results section.

### 4.4 Nonlinear embedding via *t*-SNE

To qualitatively investigate the organization of the input feature space, we perform a descriptive analysis based on *t*-distributed stochastic neighbor embedding (*t*-SNE) [41]. Specifically, we apply *t*-SNE to the input feature representations provided to the network, denoted by 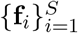, where *S* is the total number of samples and **f**_*i*_ *∈* ℝ^*N*^ . The resulting two-dimensional representation is then used for visualization purposes. In the original feature space, local relationships among samples are characterized through conditional probabilities that quantify the likelihood of selecting sample **f**_*j*_ as a neighbor of sample **f**_*i*_. These probabilities are defined as

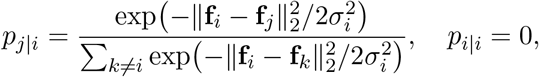

where *σ*_*i*_ denotes the bandwidth of the Gaussian kernel centered at **f**_*i*_. Consequently, samples that are closer in the original feature space are assigned higher probabilities than more distant ones. The value of *σ*_*i*_ is selected such that the entropy of the induced conditional distribution *P* _*i*_ matches a prescribed perplexity, which controls the effective number of local neighbors considered around each point. In our setting, the perplexity is set equal to the number of vectors within each semantic category, ensuring that the effective neighborhood size in *t*-SNE matches the intrinsic cardinality of each category in the dataset.

The pairwise conditional probabilities are then symmetrized to obtain joint probability coefficients

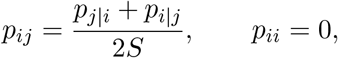

which quantify pairwise similarities among samples in the original feature space.

The *t*-SNE algorithm then seeks a low-dimensional representation of the original feature vectors. In the two-dimensional embedding space produced by *t*-SNE, each sample **f**_*i*_ is represented by a point **y**_*i*_ *∈* ℝ^2^. Pairwise similarities in this low-dimensional space are modeled through the coefficients

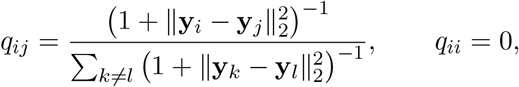

which are obtained from a Student’s *t*-distribution with one degree of freedom.

The objective of *t*-SNE is to construct a low-dimensional representation whose similarity structure, described by the coefficients *q*_*ij*_, preserves as closely as possible the similarity relationships encoded by the coefficients *p*_*ij*_ in the original feature space. The embedding is therefore computed by minimizing the Kullback–Leibler divergence [22] between the two sets of similarity coefficients:

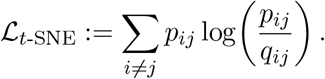

### 4.5 Learning phase

The synaptic dynamics in the network is governed by a triplet-based spike timing dependent plasticity model [32]. Let *x*_*j*_(*t*) denote the spike train emitted by the presynaptic neuron *j*. A low-pass filtered trace of the presynaptic activity is defined as

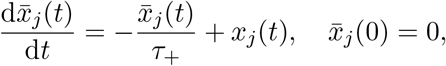

with time constant *τ*_+_ *∈* [10, 20] ms.

Similarly, the postsynaptic spike train *y*_*i*_(*t*) represents the sequence of spikes generated by the postsynaptic neuron *i* in response to the total synaptic input received from all presynaptic neurons. Two additional low-pass filtered traces were defined to capture distinct temporal dependencies in postsynaptic activity, i. e.,

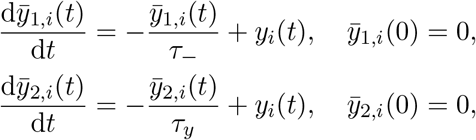

with time constants *τ*_*−*_ *∈* [20, 40] ms and *τ*_*y*_ [40, 150] ms.

The synaptic weight *w*_*ij*_(*t*) connecting the presynaptic neuron *j* to the postsynaptic neuron *i* evolves according to the minimal triplet-STDP rule [16] as

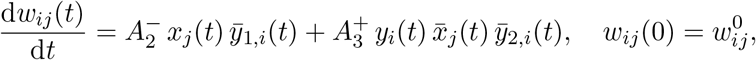

explicitly linking presynaptic firing activity *x*_*j*_(*t*) to postsynaptic responses *y*_*i*_(*t*) through temporally filtered traces that encode spike timing interactions. In particular, weight changes depend on the relative spike timing 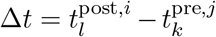, with causal ordering (Δ*t >* 0) leading to potentiation and reverse ordering (Δ*t <* 0) leading to depression.

The initial synaptic strength 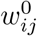 is randomly drawn from a uniform distribution in [0, 0.25], supporting synaptic competition and the emergence of functionally relevant connections [1, 37]. Consistent with [16], parameters are chosen as 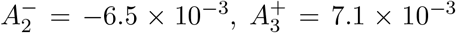, *τ*_+_ = 16.8 ms, *τ*_*−*_ = 33.7 ms, and *τ*_*y*_ = 144 ms.

Excitatory synapses are constrained to *w*_*ij*_ *∈* [0, 0.5], whereas inhibitory synapses are restricted to *w*_*ij*_ *∈* [*−*0.5, 0]. These bounds prevent unstable growth and ensure biologically plausible synaptic strengths, consistent with homeostatic regulatory mechanisms limiting excessive excitation [32].

During training, each input vector, either SFVs for the experiments reported in the Results section or DINO embeddings for those described in the DINO feature embeddings appendix, is presented for a duration of *T*_train_ = 80 ms, with no interstimulus interval. To prevent premature representational specialization and to ensure balanced exposure across hierarchical feature levels, the presentation order of feature vectors is constructed using a systematic interleaving scheme. Let **F** *∈* ℝ^*S×N*^ denote the feature matrix. As described in the External input current subsection, the rows of **F** are partitioned into two high-level meta-categories, defined by the index sets ℐ_1_ := {1, …, *S/*2} and ℐ_2_ := {*S/*2 + 1, …, *S* } (assuming for simplicity that *S* is even). Within each meta-category, features are further organized into *N*_cat_ = 5 consecutive categories, each comprising *N*_item_ = 10 feature vectors. The presentation sequence is constructed to enforce strict alternation between meta-categories while progressing systematically through categories. For a fixed item index *j* = 1, …, *N*_item_, for each category index *k* = 1, …, *N*_cat_, the *j*-th item of the *k*-th category in ℐ_1_ is immediately followed by the corresponding *j*-th item with the same category index in ℐ_2_. After all categories have been presented for a given item index *j*, the sequence advances to the item index *j* + 1, preserving the same meta-category/category alternation scheme.

### 4.6 Statistical tests

Statistical analyses are performed to assess differences between the primed and control conditions at both the level of neural firing rates and the temporal dynamics of population activation. Depending on the structure of the data and the underlying inferential question, either non-parametric rank-based tests or permutation-based cluster statistics are employed. To ensure reproducibility, the random seed 42 is fixed in all our analyses.

#### Permutation cluster test

To compare the mean firing rate of the neural network between the two conditions, while accounting for temporal dependencies across samples, we employ a non-parametric permutation cluster test [24] using the mne.stats library [18] in Python.

Let us choose a grid on the time interval [0, *T*] given by (*t*_0_, …, *t*_*M*_) with

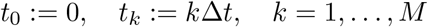

with the choice Δ*t* = 0.1 ms and *T* = *t*_*M*_ . Let (*X*_p_(*t*_0_), …, *X*_p_(*t*_*M*_)) and (*X*_c_(*t*_0_), …, *X*_c_(*t*_*M*_)) denote the mean firing rate time series of the network under the primed and control conditions, respectively. The test is applied to the pair (*X*_p_(*t*_*i*_), *X*_c_(*t*_*i*_)) for *i* = 0, …, *M* using each a two-sided statistics. At each time point, a *t*-value is computed by comparing the mean firing rates of the two conditions, producing a time-resolved series of test statistics that serves as the basis for cluster formation. Clusters are then defined as contiguous time points for which the *t*-values exceed a threshold corresponding to a two-sided significance level of *p* = 0.05. The cluster-level statistics are computed as the sum of the *t*-values within each identified cluster. Statistical significance is assessed via a permutation procedure in which condition labels are randomly reassigned across observations, thereby generating surrogate datasets under the null hypothesis of no difference between conditions. A total of *N*_perm_ = 1000 permutations are performed on the entire time series. For each permutation, the cluster-level statistics are computed for all clusters, and only the maximum value is retained to form the empirical null distribution. This procedure controls the family-wise error rate, i. e., the probability of observing at least one false positive across all clusters, ensuring that the overall significance level is maintained. Observed clusters are considered statistically significant if their associated *p*-value, obtained by comparison with this null distribution, is below the significance threshold of 0.05.

#### Wilcoxon signed-rank test

To identify neurons exhibiting a statistically significant stimulus-evoked response, we employ the Wilcoxon signed-rank test [43], a non-parametric alternative to the paired-samples *t*-test. The test is implemented in Python using the pingouin library [40].For each neuron and each stimulus with at least 10 paired reference–response observations, we compare trial-by-trial mean firing rates measured during the current 300 ms stimulus-presentation window with those measured during the immediately preceding 300 ms stimulus-presentation window. Because no preceding stimulus is available for the first presentation of each sequence, this analysis is restricted to the second through fifth presentations. The test is conducted in a two-sided manner, with the null hypothesis that response and reference firing rates do not differ. A neuron is provisionally classified as responsive if at least one stimulus yields a *p*-value *<* 0.05. This step serves as an initial screening criterion, neurons passing this criterion subsequently undergo additional selection steps based on their stimulus-specific Z-scores, described in the Results section, before being included in the final rank-based classification analyses.

#### Mann-Whitney *U* –test

To test for differences in the temporal onset of neural population activation between the two conditions, we employ the Mann-Whitney *U* -test [29], a non-parametric alternative to the independent-samples *t*-test. The test is implemented using the pingouin library [40] in Python. The Mann-Whitney *U* -test evaluates whether two samples are drawn from the same distribution by comparing their rank orderings, without assuming normality or homoscedasticity. All tests are conducted in a two-sided manner, with the null hypothesis that the distributions of activation times in the two conditions do not differ. To account for multiple comparisons where applicable, *p*-values are corrected using the Holm-Bonferroni method [2], controlling the family-wise error.

## 5 Simulation environment

All simulations are performed on a high-performance workstation. The system is equipped with an AMD Ryzen Threadripper PRO 7975WX CPU (64 cores, up to 5.35 GHz), 128 GB of RAM, and an NVIDIA RTX 6000 Ada Generation GPU with 48 GB of VRAM. The computations are executed on Ubuntu Linux 24.04.3 LTS with the 6.8.0-79-generic kernel.

## 6 Code availability

The source code is available at https://github.com/didibar/neural-adaptation-adex-triplet-stdp.git.

## 7 Acknowledgements

We thank Janko Petkovic and Lorenzo Squadrani from the University of Bonn for their valuable insights and stimulating discussions.

## 8 Author Contributions

**Diletta Bartolini**: Methodology, Software, Validation, Formal analysis, Writing - Original Draft, Writing - Review & Editing; **Thomas P. Reber**: Conceptualization, Data Curation, Writing - Review & Editing, Supervision; **Tatjana Tchumatchenko**: Writing - Review & Editing, Supervision; **Matthias Voigt**: Conceptualization, Formal analysis, Writing - Review & Editing, Supervision, Funding acquisition. All authors contributed to commenting on the manuscript.

## 10 Additional information

Correspondence and requests for material should be addressed to Diletta Bartolini.

## A DINO feature embeddings

In this *Appendix*, we present a comparative analysis of the networks behavior when using DINO feature embeddings (FVs) versus synthetic feature vectors, whose characteristics and performance are extensively discussed in the Results section.

The synthetic and DINO feature spaces exhibit fundamentally different organizations. Synthetic features are characterized by larger Euclidean distances, particularly between categories, resulting in a more segregated and pronounced categorical structure. In contrast, DINO features show smaller variations, leading to a more compact distribution, as depicted in Figure 7.

This contrast is further reflected in the corresponding low-dimensional *t*-SNE embeddings shown in Figure 8. For SFVs, the representation exhibits a well-defined hierarchical organization of the dataset, as discussed in the Results section. In contrast, the DINO feature space shows a weaker degree of categorical separation. While local neighborhood relationships are partially preserved, neither a clear partition between the two principal meta-categories nor a consistent subdivision into categories emerges. The resulting embedding is more diffuse, with limited cluster compactness and a small number of isolated points.

At the population level, as shown in Figure 9, neuronal responses to DINO FVs exhibit comparable firing rates between primed and control conditions at stimulus onset. Throughout the response period, the primed condition shows reduced peak firing rates relative to the control condition, with a single time window displaying a statistically significant difference (cluster-based permutation test, significant time window: [925.6, 926.0] ms, cluster size = 5, sum of *t*-values = *−*21.8, *p* = 2.4 *×* 10^*−*2^). Over extended timescales and in the absence of additional sensory input, network activity gradually returns to baseline, following a comparable decay trajectory and converging toward similar average firing rates in both conditions. Detailed firing-rate time courses are provided in *Table_SourceData_DINO_FVs*.*xlsx* file, available from the repository linked in Code availability.

**Figure 9:**
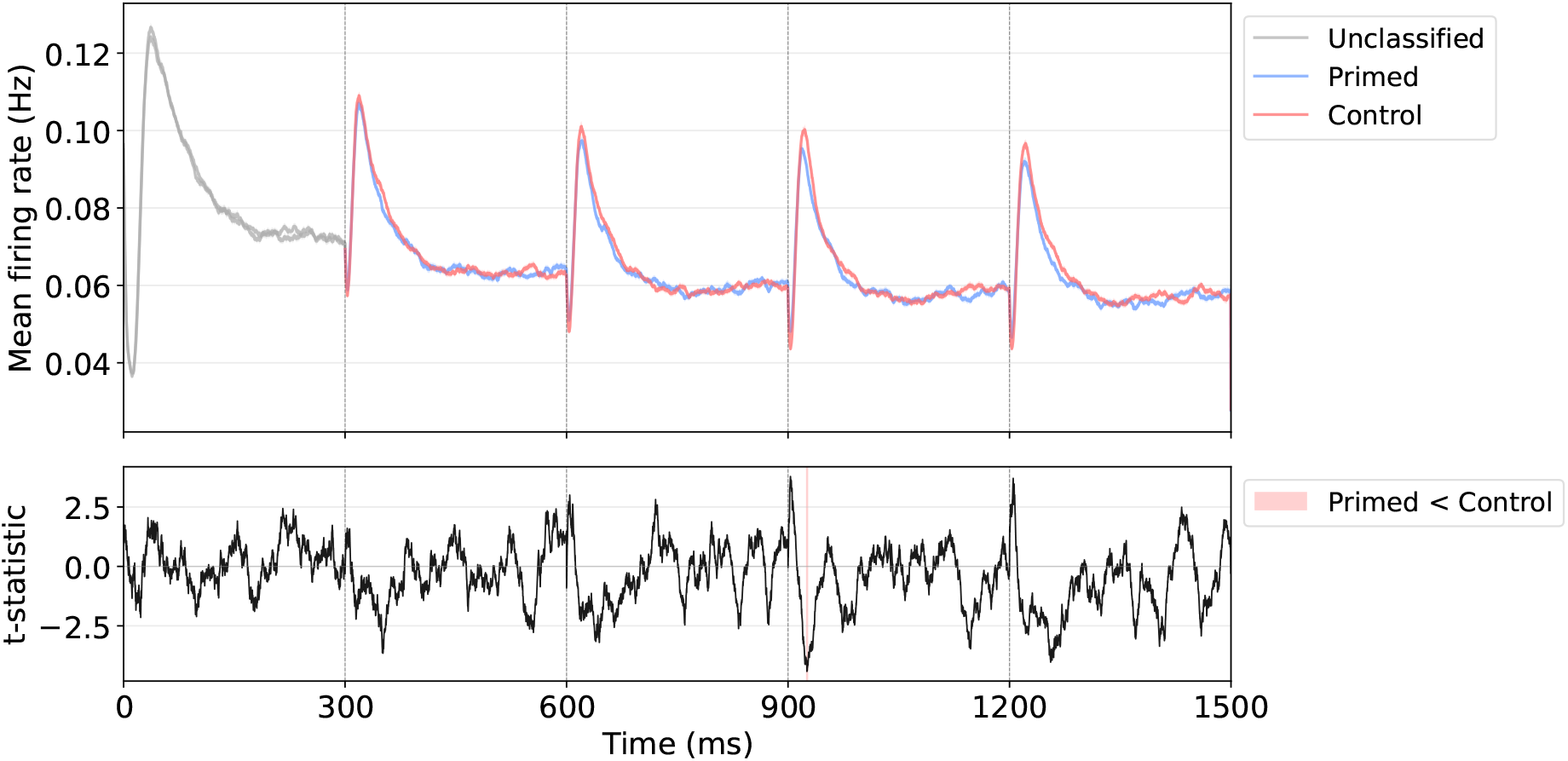
Temporal evolution of network activity. The figure shows the temporal evolution of mean firing rates for primed and control conditions and the corresponding cluster-based statistical comparison using DINO feature embeddings as input. All conventions are identical to those described for the SFV analysis in Figure 2.

Regarding the dynamics of neuronal recruitment, the temporal patterns observed using DINO feature embeddings are qualitatively consistent with those obtained with SFVs. At the 10% activation level, neurons under the primed condition appear to reach the spiking threshold earlier than those under the control condition (Figure 10a). At the 15% level, a reversed pattern is observed, with shorter latencies in the control condition (Figure 10b). However, neither effect reaches statistical significance (Table 8 and Table 9). Similarly, the single-neuron onset latency analysis (Figure 11) does not reveal any statistically significant trend (Table 10).

**Figure 10:**
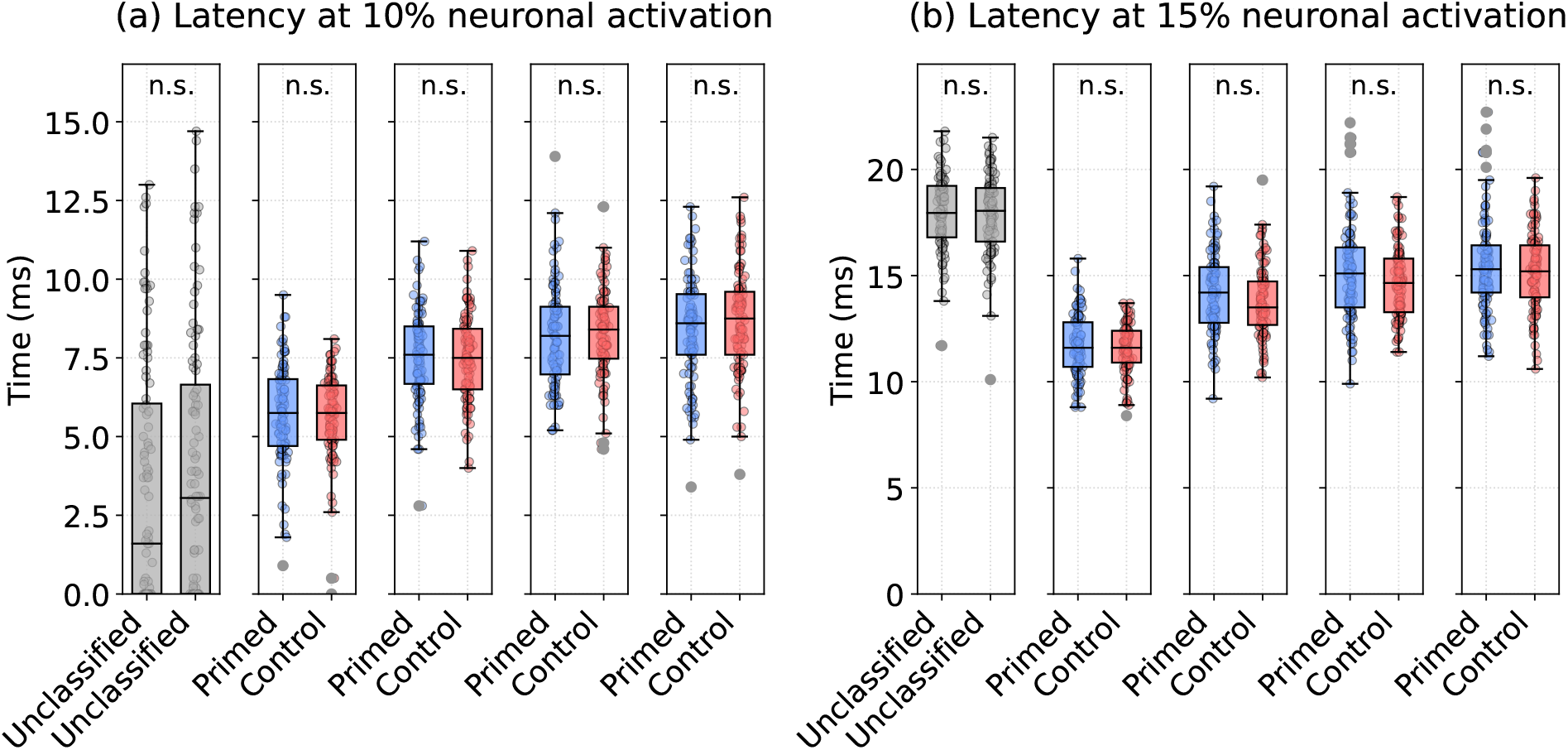
Neuronal recruitment dynamics. Same analysis as in Figure 3, using DINO feature embeddings. Box plots show the distribution of activation latencies for (a) 10% and (b) 15% neuronal recruitment thresholds. Primed and control conditions are shown in blue and red, respectively, while the first time window (gray) is not assigned to either condition.

**Table 8:** Neuronal activation latency at the 10% threshold across time windows using DINO feature embeddings. Statistical comparisons between conditions were performed using the Mann–Whitney *U* -test with Holm correction for multiple comparisons. Same analysis as in Table 3.

| Time window (ms) | primed | control | $U$ | $p$-value |
| --- | --- | --- | --- | --- |
|  | median [Q1, Q3] | median [Q1, Q3] |  |  |
| [300, 600] | 5.8 [4.7, 6.8] | 5.8 [4.9, 6.6] | 5100.0 | 1.0 |
| [600, 900] | 7.6 [6.7, 8.5] | 7.5 [6.5, 8.4] | 5086.0 | 1.0 |
| [900, 1200] | 8.2 [7.0, 9.1] | 8.4 [7.5, 9.1] | 4645.0 | 1.0 |
| [1200, 1500] | 8.6 [7.6, 9.5] | 8.8 [7.6, 9.6] | 4586.0 | 1.0 |

**Table 9:**
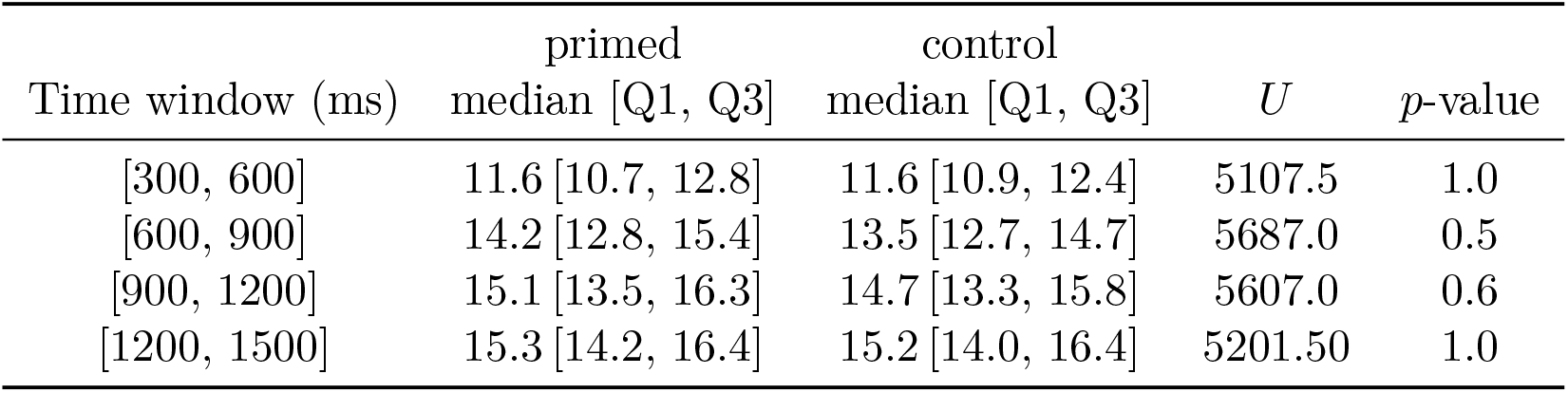
Neuronal activation latency at the 15% threshold across time windows using DINO feature embeddings. Reporting format and statistical procedure are as described in Table 3.

| Time window (ms) | primed | control | $U$ | $p$-value |
| --- | --- | --- | --- | --- |
|  | median [Q1, Q3] | median [Q1, Q3] |  |  |
| [300, 600] | 11.6 [10.7, 12.8] | 11.6 [10.9, 12.4] | 5107.5 | 1.0 |
| [600, 900] | 14.2 [12.8, 15.4] | 13.5 [12.7, 14.7] | 5687.0 | 0.5 |
| [900, 1200] | 15.1 [13.5, 16.3] | 14.7 [13.3, 15.8] | 5607.0 | 0.6 |
| [1200, 1500] | 15.3 [14.2, 16.4] | 15.2 [14.0, 16.4] | 5201.50 | 1.0 |

**Figure 11:**
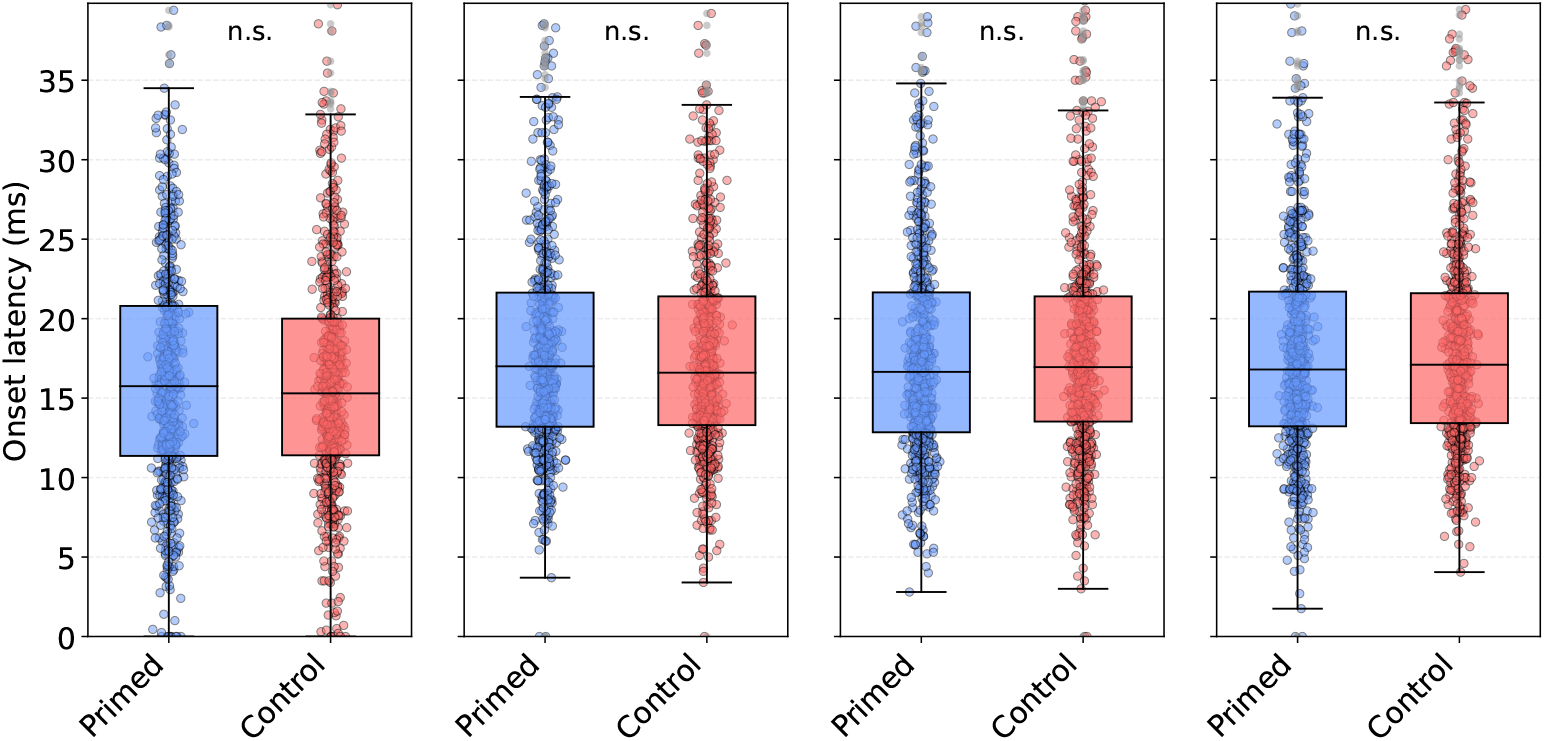
Single-neuron onset latency. Same analysis as in Figure 4, but using DINO feature embeddings as input. Box plots show the distribution of single-neuron onset latencies under primed (blue) and control (red) conditions across time windows. In the first time window (gray), conditions are not yet differentiated. Statistical procedures follow those described for the SFV analysis.

**Table 10:** Single-neuron onset latency. Same analysis and reporting as in Table 5, but using DINO feature embeddings as input. Statistics across time windows are computed under the same inclusion criterion and statistical significance is assessed using a two-tailed paired permutation test with Holm correction for multiple comparisons across time windows.

| Time window (ms) | $n$ | primed | control | $\Delta_{\text{median}}$ | $p$-value |
| --- | --- | --- | --- | --- | --- |
|  |  | median [Q1,Q3] | median [Q1,Q3] |  |  |
| [300, 600] | 730 | 15.8 [11.0, 20.5] | 15.3 [11.0, 19.6] | 0.1 | 1.0 |
| [600, 900] | 730 | 17.0 [12.8, 21.2] | 16.6 [12.6, 20.7] | 0.0 | 1.0 |
| [900, 1200] | 731 | 16.6 [12.2, 21.0] | 17.0 [13.0, 20.9] | -0.1 | 1.0 |
| [1200, 1500] | 727 | 16.8 [12.6, 21.0] | 17.1 [13.0, 21.2] | -0.1 | 1.0 |

At the population level, DINO-based results are broadly consistent with the trends reported in the Results section, however, the distinction between primed and control conditions is less pronounced, and statistical evidence is weaker compared to the synthetic input case.

At the single-neuron level, applying the same analysis to DINO feature embeddings does not reveal a consistent stimulus-driven structure across neurons, and no robust functional segregation is observed.

Overall, DINO-based results are qualitatively consistent with the population-level trends reported in the Results section, but the distinction between primed and control conditions is substantially weaker and lacks statistical significance. This is expected from the geometric analysis: the less structured DINO feature space corresponds to a low-*α*, low-*β* operating regime, in which our parameter sweep predicts reduced sharpening expression and shorter suppression windows.

## Footnotes

1 Note that the triplet-based spike timing dependent plasticity rule is an abstract model, meaning that all quantities in the model are dimensionless. However, in order to use it in (2), the synaptic input provided by this model must be a conductance. For this reason, we explicitly added the unit nS in (3) and (4).

